# Molecular flux control encodes distinct cytoskeletal responses by specifying SRC signaling pathway usage

**DOI:** 10.1101/648030

**Authors:** Adèle Kerjouan, Cyril Boyault, Christiane Oddou, Edwige Hiriart-Bryant, Alexei Grichine, Alexandra Kraut, Mylène Pezet, Martial Balland, Eva Faurobert, Isabelle Bonnet, Yohann Coute, Bertrand Fourcade, Corinne Albiges-Rizo, Olivier Destaing

**Affiliations:** Institute for Advanced Biosciences, Centre de Recherche Université Grenoble Alpes, Inserm U 1209, CNRS UMR 5309, France; Laboratoire EDYP, BIG-BGE, CEA Grenoble France; Laboratoire interdisciplinaire de physique (Liphy), Univ. Grenoble Alpes, CNRS, 38000, Grenoble, France; Laboratoire Physico-Chimie Curie, Institut Curie, PSL Research University, Sorbonne university, UMR 168, 75005, Paris, France

**Author notes:** To whom correspondence should be addressed, **Institute for Advanced Biosciences, Centre de Recherche UGA / Inserm U 1209 / CNRS UMR 5309, Site Santé - Allée des Alpes 38700 La Tronche**, France.

**Keywords:** SRC, conformational intermediates, optogenetics, time-resolved phospho-proteomics, pleiotropy, migration, invasion, encoding intracellular signaling, cytoskeletal, adhesions

## Abstract

Multi-domain signaling proteins sample numerous stimuli to coordinate distinct cellular responses. Understanding the mechanisms of their pleiotropic signaling activity requires to directly manipulate their activity of decision leading to distinct cellular responses. We developed an optogenetic probe, optoSRC, to control spatio-temporally the SRC kinase, a representative example of versatile signaling node, and challenge its ability to generate different cellular responses. Genesis of different local molecular fluxes of the same optoSRC to adhesion sites, was sufficient to trigger distinct and specific acto-adhesive responses. Collectively, this study reveals how hijacking the pleiotropy of SRC signaling by modulating in space and time subcellular molecular fluxes of active SRC kinases.

## Introduction

The numerous extracellular and intracellular stimuli are integrated by a limited number of multi-domain signaling molecules that coordinate cell adaptation and behaviors. How specific cellular responses are generated through precise control of the spatiotemporal dynamics of activation of one signaling molecule has remained a fundamental open question.

To investigate this paradigm, we focused on the first discovered proto-oncogene, c-SRC, which is the prototype of the SRC family kinases (SFKs). Each member of this family is highly pleiotropic since it regulates diverse cellular outputs, such as metabolism, proliferation, gene expression and cell morphology^1^. The ability of c-SRC signaling to decide between distinct cellular responses regulating cell-substrate and cytoskeletal remodeling clearly sustained its role of decision-making in numerous migration and invasion processes^2^. C-SRC can even further displays antagonistic functions in acto-adhesive structures as invadosomes, where it regulates both assembly and dismantling of these structures^3^. Consequently, decrypting the numerous potential c-SRC signaling activities in the context of these dynamic acto-adhesive structures is challenging, as these activities occur at micrometer and minute scales.

To investigate the mechanism of SRC encoding activity, we need to integrate all structural elements that allow c-SRC to sample inputs, to route specific signaling pathways among a large repertoire and to encode specific cellular phenotypes. All SFK members share a canonical structure in which their kinase domain is targeted to membranes by a SH4 domain and maintained fully closed by two intramolecular interactions: one between the SH2 domain and a C-terminal phosphorylated tyrosine and the second between the SH3 domain and an internal proline rich region (PRR)^4^ Release of these intra-molecular interactions fully activates c-SRC kinase activity, through the opening of its bi-lobal kinase domain and provoking a cascade of events: autophosphorylation of the Y416 residue ^5^ and solvent exposure of the adaptor SH3, PRR and SH2 domains, which become further available for extra-molecular interactions. Therefore, the control of the conformational changes of SRC allows a tight coupling of its enzymatic activation with its adaptor functions.

According to the structural complexity of c-SRC, one can predict that c-SRC signaling is not based on a simple binary activation but instead relies heavily on the relationship between a multitude of possible c-SRC conformational intermediates defined by inter-domain interaction combinations (Fig. 1A, B). In agreement with this hypothesis, it has been reported that its kinase activities and adaptor functions are affected at various levels by conformational changes of c-SRC^5^. First, its kinase activity is highly regulated by the precise positioning of the SH2 and SH3 domains^4^. Any perturbation of the two intramolecular regulatory interactions by dephosphorylation of the regulatory C-ter tyrosine or binding of any SRC domains (unique domain-UD, SH3, PRR, SH2 domains or phosphorylated regulatory C-ter tyrosine) with extra-molecular domains (Fig. 1B) is known to modulate the duration and level of SRC kinase activity^6^. Second, interactions with lipids add an additional layer of complexity by affecting the binding capacities of the poorly structured UD and SH3 domains^7^. Third, this complexity also includes the level of clustering of this kinase while being able to oligomerized from the poorly knowns c-SRC dimers to 80 nm-nanoclusters of c-SRC molecules ^8–10^. The vast amount of SRC conformational intermediates suggests that c-SRC signaling emerges from a heterogeneous population of molecules with different levels of kinase activity targeting dynamically a repertoire of interactors. Understanding the functional relevance of the multiple c-SRC conformational intermediates in a dynamical manner is impossible with classical genetic methods (Fig. 1A). Optogenetics manipulations have emerged as a powerful tool to dynamically regulate cell signaling^11^. Nevertheless, with a temporal resolution ranging from 3 to 300s and a spatial resolution superior to 5 μm^12, 13^, it remains challenging to access the higher spatial resolution necessary to control acto-adhesive structures. Chemogenetics has been used to directly targeted the selectivity of SRC lacked both minute-scale and reversible control of SRC at the subcellular level^14, 15^, while optogenetics was only used to obtain reversible and subcellular photo-inhibition of SRC signaling^16^.

**Figure 1.**
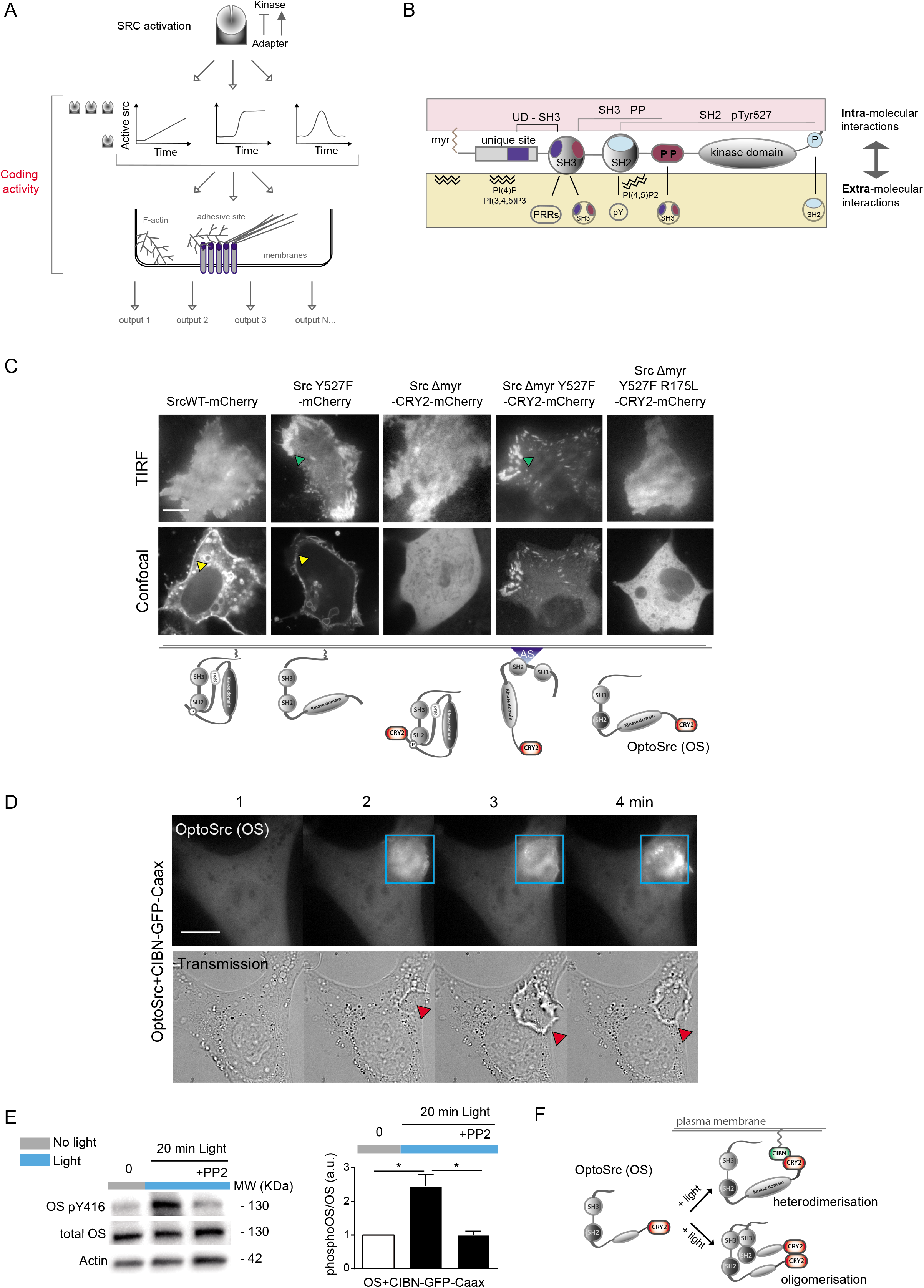
Optogenetic control of the non-receptor tyrosine kinase SRC. **A.** Schematic of how the regulatory coupling between adaptor and kinase activities of SRC could trigger different cellular outputs by sustaining a population of numerous SRC conformational intermediates. **B.** High combinatorial of SRC conformations is sustained by numerous intra- and extra-molecular interactions. **C.** Representative images of different optogenetic probes engineered and expressed in MDCK cells maintained in the dark, that led to the optoSrc (OS) which is a potentially active tyrosine kinase fully cytosolic while not localized in ECM-adhesive sites (blue arrows) and cell-cell contacts (yellow arrows). **D.** Representative time serie showing that local recruitment of OS (blue square) at the plasma membrane of fibroblasts expressing CIBN-GFP-Caax triggers large dorsal ruffles (red arrows). **E.** Light-dependent recruitment of OS at the plasma membrane of MDCK cells expressing CIBN-GFP-Caax induces SRC-dependent activation of its kinase domain (measured by P416Y/total SRC ratio). **F.** OS photostimulation can lead to either its membrane relocalization (through heterodimerization with membrane-anchored CIBN) or its oligomerization. Scale bars: 5 μm.

To spatio-temporally activate and control SRC signaling, we engineered a new photosensitive SRC kinase, optoSRC (OS) based on the CRY2 optogenetic module. We showed that sustained light-dependent OS oligomerization was sufficient to induce its SH3-dependent relocalization to adhesive sites by diffusive transport and, then, create there a subcellular localized and sustained flux of OS. We were able to modulate flux of OS oligomers to induce different acto-adhesive phenotypes or regulate the intensity of these cellular responses. Finally, this synthetic approach was sufficient to hijack the fundamental principles of SRC coding activity by demonstrating the causal link between the generation of specific fluxes of OS oligomers and the dynamics of downstream signaling pathways to trigger different cellular responses.

## Results

### Development of a light-inducible SRC kinase (optoSRC, OS)

Previous biochemical reports^5^ have highlighted that the large repertoire of possible c-SRC conformations that supports its signaling complexity through the coupled dynamics of its kinase activity and adaptor functions (Fig. 1A). According to numerous possible intra- or extra-molecular interactions (Fig. 1B), it is not clear how spatio-temporal control of c-SRC activity trigger different cellular ouputs (Fig. 1A, B). Precise spatial control of c-SRC activation is especially important for acto-adhesive structures, such as lamellipodia or invadosome rings (Fig. S1A).

Besides trying to reproduce the complexity of endogenous activation of SRC, we developed a CRY2-dependent optogenetic system^17^ to spatio-temporally control its activity based on reported SRC mutations that both affects its membrane localization and the opening of its kinase domain. Our strategy was to create a potentially active and cytosolic SRC mutant displaying only SRC signaling events upon light stimulation while being maintained inactive in the dark. To do so, the CRY2-mCherry module was fused to a SRC mutant deleted of its membrane-anchoring domain (SRCΔmyr-CRY2-mCherry), leading to its absence of localization in cell-cell contacts or adhesive sites of MDCK cells differently from c-SRCWT-mCherry or the active mutant SRCY527F-mCherry (Fig. 1C). This mutant was then made potentially active by opening its closed conformation through the substitution of phenylalanine at regulatory Y527. Despite not accumulating at cell-cell contacts, the SRCΔmyr,Y527F-CRY2-mCherry mutant was able to localize to adhesive sites in the dark (Fig.1C). An additional point mutation, R175L, inhibited the PY-binding activity of the SH2 domain and abolished the ability of SRCΔmyr,Y527F-CRY2-mCherry to localize in adhesive sites. This final construct named optoSRC (OS; SRCΔmyr,R175L,Y527F-CRY2-mCherry mutant) did not accumulate in any cellular compartment in the dark (Fig. 1C).

We next validated the functionality of OS by assessing the induction of a typical SRC-driven cellular response by light upon its membrane relocalization. OS relocalization was obtained by CRY2 heterodimerization with a plasma membrane anchored CIBN (CIBN-GFP-Caax) in response to local light stimulation (Fig. 1D). As expected for CRY2-CIBN heterodimerization, sustaining cyclic light stimulation induced periodic OS membrane localization with a 5 μm-spatial resolution (Fig. S1B). Local membrane recruitment of OS provoked the rapid formation of large, localized dorsal ruffles (Fig.1D, Suppl. Video 1) and phenocopied the expression of a constitutively active mutant of SRC (SRCY527F, Fig. S1C), the activation of a thermosensitive SRC mutant^18^ or a chemo-activable SRC^19^. The light-dependent membrane recruitment of OS was associated with self-activation of its kinase domain, as indicated by PP2-sensitive auto-phosphorylation of its Y416 residue (Fig. 1E). Furthermore, the specific light-dependent activation of OS and the low leakiness of OS in the dark was confirmed by analyzing paxillin (PXN) phosphorylation, the main cytosolic substrate of c-SRC (Fig. S1D).

Altogether, our data show that OS was a fully functional optogenetic SRC system mimicking physiological SRC functions.

### OS nanoclustering drives its relocalization to adhesive sites

Along with its light-activable kinase, OS also reduced intra-molecular interactions, thereby decreasing the combination of potential SRC conformational intermediates. In addition to OS and CIBN heterodimerization, the ability of CRY2 to oligomerize was used to investigate the properties of other SRC intermediates (Fig. 1F) such as SRC dimers or SRC nanoclusters, whose *in vivo* functions are poorly characterized. To characterize the oligomerization level of light-activated OS in our experimental conditions, we measured the variations of OS molecular brightness as it is proportional to the oligomerization state of a fluorophore^20^, with a live imaging method based on FCS measurements. In response to blue light, OS presented a significant increase of brightness that became comparable to the brightness of two OS that were molecularly fused (double OS) and maintained in the dark (Fig.S2A). This supports that activated OS can form small oligomers (from dimers to low oligomers) and that light-activation of OS alone is founded to study the influence of SRC nanoclusters (small oligomers) on its signaling functions.

We used blue light TIRF illumination to induce mostly OS oligomers at the vicinity of the plasma membrane. Surprisingly, formation of OS oligomers induced their rapid and specific relocalization from the cytosol to adhesive sites, as they colocalized with the focal adhesion markers vinculin, p130CAS and paxillin (Fig. 2A-B; Fig. S2D; Suppl. Video 2) but not with clathrin or caveolin sites, which are other c-SRC-sensitive plasma membrane-targeted loci (Fig. S2B). This relocalization was not dependent on oligomerization with endogenous SFK members as it was still effective in SYF cells (*c-Src^-/-^ Yes^-/-^ Fyn^-/-^* cells; Fig. 2C). Light-dependent OS relocalization to adhesive sites was also typical of the kinetics of CRY2 activation (Fig. S2E) and exhibited poor leakage of its phosphorylation activity under dark conditions, as illustrated by paxillin phosphorylation in MDCK cells or SYF cells (Fig. S3A, B).

**Figure 2:**
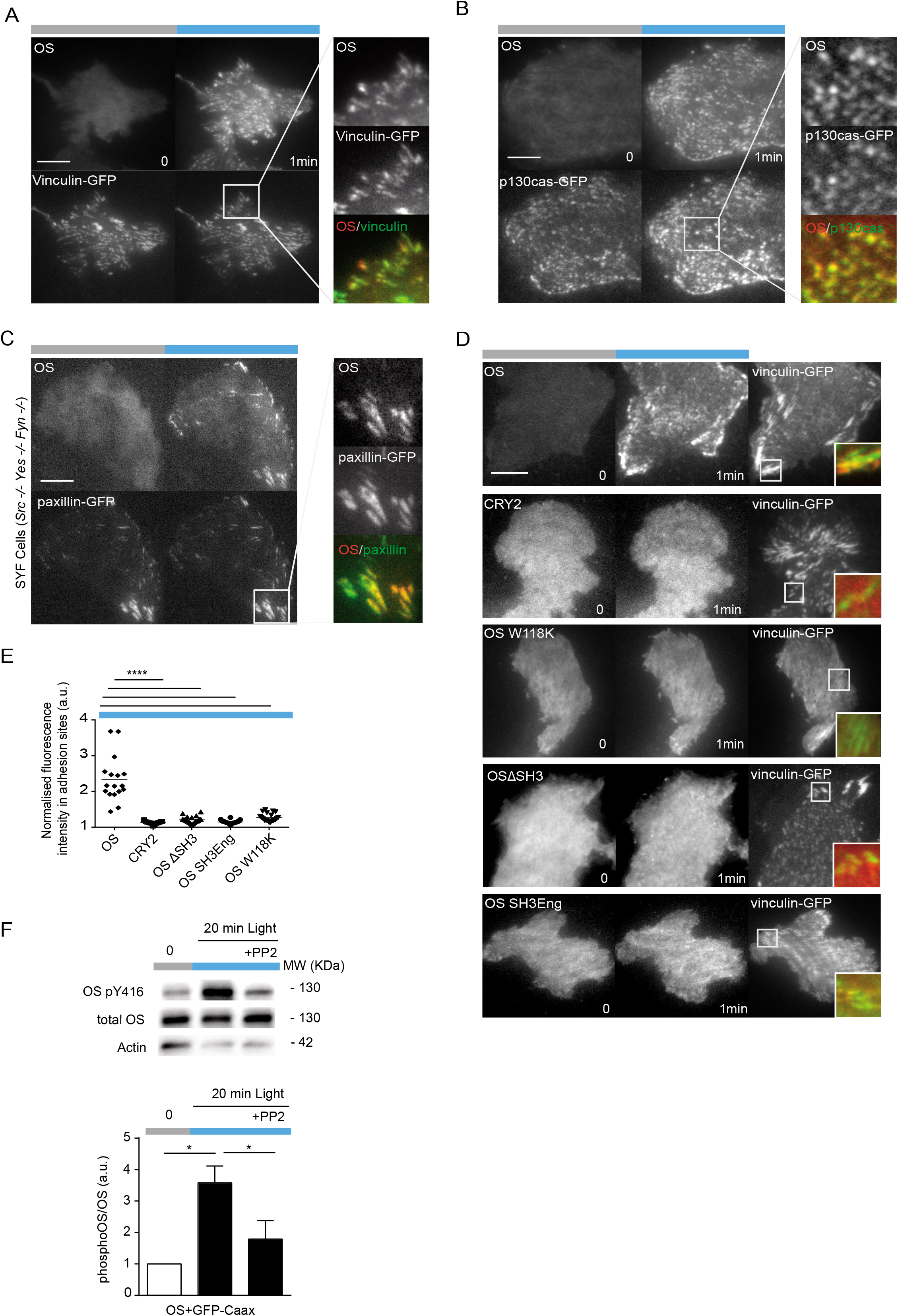
Light-dependent OS oligomers relocalize to adhesive sites through their SH3 domains. **A-B.** Representative TIRF images before and 1 min after blue TIRF photostimulation showed relocalization of OS oligomers to adhesive sites of MDCK cells since colocalizing with vinculin-GFP (B) or p130cas-GFP (C). **C.** Representative TIRF image showed that relocalization of OS oligomers in adhesive sites (paxillin-GFP) even occurs in SYF cells and thus in absence of the main endogenous SFKs. **D.** Representative TIRF images showed that light-dependent OS oligomers relocalization in adhesive sites of MDCK cells is dependent of its SH3 domains, while not observed for CRY2 alone or the following OS mutants (OSΔSH3, OS W118K and OS SH3Eng). **E.** Quantification of OS and its SH3-mutants at the plasma membrane after blue TIRF photostimulation (SD; N=3; >30 cells per condition, unpaired t test). **F.** Light-dependent oligomerization of cytosolic OS mediates activation of its kinase domain (measured by P416Y/total SRC ratio) through a PP2-sensitive process. Scale bars: 5 μm.

Relocalization of OS oligomers to adhesive sites was supported by the SH3 domain functions (Fig. 2E and S3G) as evidenced by its strong decrease upon SH3 deletion (OS ΔSH3), inhibition of SH3 domain binding to PRR-proteins (OS W118K) or increase of intramolecular SH3-PRR binding (OS SH3Eng or SRC-K249P/Q252P/T253P^21^). In addition, mutation of the internal PRR of OS (OS PPR-AAA) had no effect on this process (Fig. S2). Interestingly, the presence of the phosphorylable Y527 in OS (OS R175L only) blocked its ability to be relocalized in adhesive sites showing that this tyrosine 527 not only controled the kinase domains through the opening of the internal SH2-Y527 bridge but was also implicated in OS partners binding when phosphorylated (Fig.S2C). However, light-dependent OS relocalization could be either driven by a simple increase of SH3-local concentration induced by the oligomerization or controlled by a specific SRC-SH3 domain conformation which can be dictated by the vicinity of the SH2 domain and/or tyrosine kinase domain. This last hypothesis was confirmed by the reduced ability of activated OS mutants containing a deleted kinase domain or SH2 domain, or perturbation of the kinase domain (K295M) to relocalize in adhesive sites (Fig. S2C, D). Finally, reducing clustering properties of CRY2 by using CRY2 low mutant (CRY2-1 to 488-EED,^22^) slightly reduced OS relocalization in adhesive sites without blocking it (Fig.S2C) showing that lower-size oligomers (probably dimers) can support this relocalization.

In addition to regulate OS adaptor functions, we investigated whether SRC nanoclustering also activated its kinase activity. Light-dependent oligomerization of OS was sufficient to stimulate itself OS kinase activation, as indicated by the increase of Y416 phosphorylation (Fig. 2F), as it was PP2-sensitive (Fig. 2F) and also occurring in SYF cells (Fig. S3B). Then, OS kinase activation was increased by a light-dependent trans-phosphorylation process. In SYF cells, light-induced oligomerization of OS activated its kinase domain and phosphorylate paxillin at the same level than c-SRC re-expression (Fig. S3B).

Controlling oligomerization formation uncovered the crucial role of the SH3 domain as another mechanism for SRC nanocluster to regulate kinase activation and sample environment by mediating SRC stabilization in adhesive sites. Light-dependent OS oligomers subcellular relocalization in adhesive sites demonstrated therefore marked improvement of the spatial resolution of previous CRY2 optogenetics approach.

### Optogenetic control of the different rates of OS recruitment to adhesive sites

Next, based on the high spatial resolution of our optogenetic approach, we investigated whether different quantities of OS could be specifically targeted to adhesive sites in order to induce distinct SRC-dependent signaling events. To this end, we modulated OS oligomers recruitment rates to adhesive sites by anchoring them on membrane through co-expression of CIBN-GFP-Caax. Indeed, CRY2 oligomerization and CRY2–CIBN heterodimerization coexist as they are supported by different regions of CRY2^22^. First, we determined if membrane-associated OS oligomers (OS + CIBN-GFP-Caax) could also relocalized to adhesive sites after sustained TIRF illumination (Fig. 3A; Suppl. Video 4). In the presence of CIBN-GFP-Caax, OS oligomers were recruited with the same rate outside (membrane) and inside of adhesive sites (Fig. 3B). By contrast, OS oligomers alone (OS + GFP-Caax) were mainly targeted to adhesive sites (Fig. 3C).

**Figure 3.**
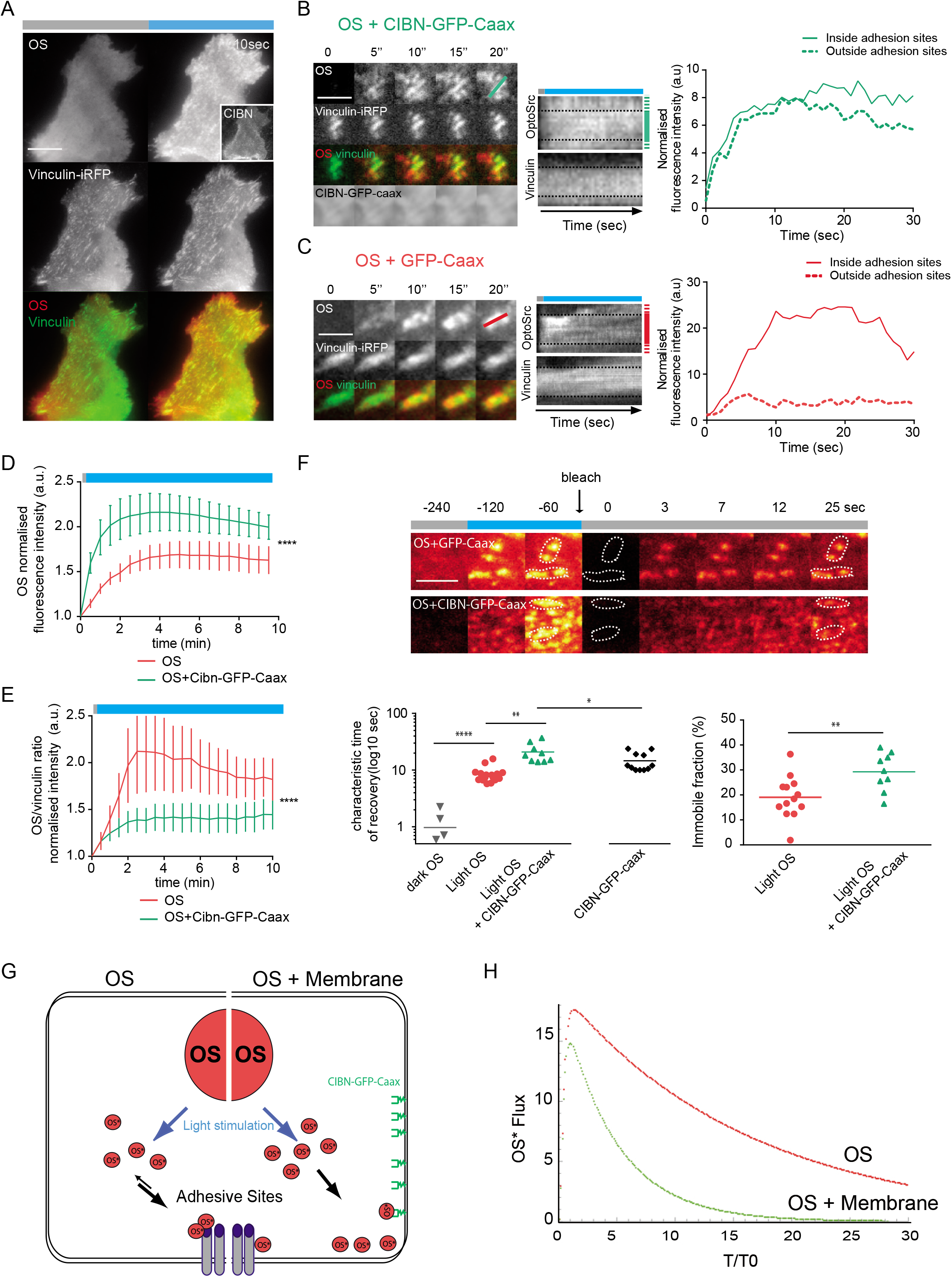
Modulation of activated OS molecular flux in adhesive sites by controlling its membrane anchoring. **A.** Representative TIRF image of light-dependent OS oligomers relocalization in adhesive sites (vinculin-iRFP) after membrane recruitment mediated by co-expression of CIBN-GFP-Caax in MDCK cells. **B-C.** Time series and kymograph characterized relocalization dynamics in response to blue TIRF photostimulation of OS oligomers inside and outside (dashed line) of adhesive sites when co-expressed in MDCK cells (B) with CIBN-GFP-Caax or (C) the control GFP-Caax. **D-E.** Quantification of OS relocalization at the whole basal level of the plasma membrane (D) or specifically in adhesive sites (F, OS/vinculin-iRFP ratio) after blue TIRF photostimulation of MDCK cells expressing either CIBN-GFP-Caax or GFP-Caax (SD; N=3; >30 cells per condition, unpaired t test) **F.** Representative TIRF time series of OS relocalization to adhesive sites (white dashed line) after blue TIRF photostimulation, followed by FRAP experiments. Quantification of FRAP parameters (SD; N=3; >15 cells per condition; unpaired t test). **G.** Physical principles of a cytosolic OS reservoir that can be relocalized to adhesive sites after its photoactivation (OS*, OS oligomers), directly or through a membrane step, mimicking respectively GFP-Caax and CIBN-GFP-Caax conditions. **H.** Numerical simulations of the mean flux of OS* per unit length of an adsorbing cluster and over time, directly (red) or through a membrane step (green). Scale bars: 5 μm (A), 2,5 μm (B, C, F).

While these two experimental strategies (direct or membrane-based indirect recruitment) both led to sustained OS recruitment to adhesive sites, we compared the rates of OS passing through these sites in terms of quantity and molecular mobility. At the whole basal cell surface, co-expression of CIBN-GFP-Caax allowed recruitment of more OS oligomers at the plasma membrane than when OS was expressed with GFP-Caax (Fig. 3D), probably because of the difference between the large amounts of available membrane anchored CIBN-GFP-Caax and the small number of adhesive sites. However, OS oligomers were more concentrated in adhesive sites than when associated with membranes (OS + CIBN-GFP-Caax), as revealed by their higher OS/vinculin-iRFP ratio after 10 min photostimulation (33 mHz, Fig. 3E). Therefore, these two strategies of OS recruitment generated different densities of OS in adhesive sites over time. In addition to determine the quantity of activated OS in adhesive sites, it was also essential to analyze the molecular mobility of activated OS in both conditions (GFP-Caax or CIBN-GFP-Caax). Thus, we combined optogenetic manipulation with FRAP analysis to determine the mobility of activated OS molecules in adhesive sites. In comparison to direct recruitment of OS oligomers to adhesive sites, membrane-based indirect recruitment (in presence of CIBN-GFP-Caax) strongly increased the immobilization of activated OS in adhesive sites (characteristic time of recovery evolving from of 8,4 s ^+^/_-_ 2,6 s to 20,8 s^+^/_-_ 8,3, associated with a 50% increase of the immobile fraction, Fig. 3F).

Therefore, we were able to generate different quantities and molecular mobilities of activated OS in adhesive sites over time by controlling its modes of recruitment.

Then, we characterized the two rates of activated OS recruitment to adhesive sites in response to sustained cyclic light activation by modeling. Our optogenetic system can be schematized as a model in which light activates an OS buffer in the cytosol. Following sustained cyclic photostimulation, this activated buffer (OS*) is flushed either by direct adsorption on the membrane (mimicking CIBN-GFP-Caax condition) or direct recruitment to adhesive sites (mimicking GFP-Caax condition) (Fig. 3G). A first consequence of this model was a 3D-2D reduction of the dimensionality of the mobility for membrane recruitment of OS* compared with direct adsorption of OS* to adhesive sites, explaining FRAP results. Indeed, we expected the rate of recovery of OS oligomers in adhesive sites to be largely smaller for a 2D diffusing membrane anchored protein (*D*≈ 0,2 μm^2^.s^-1^) than for a 3D diffusing cytosolic OS oligomers (*D*≈ 10 μm^2^.s^-1^, Fig. 3F).

Based on our experimental results, sustained cyclic light activation of OS induces a transport by diffusion of these activated oligomers (OS*) from the cytosol to adsorption sites (composed of receptors for OS*), generating a local diffusion-limited flux of activated OS controlled by the density of receptors composing the sites. To compare both OS* fluxes, numerical simulation was used to compare the early behavior of OS fluxes that were directly recruited to adhesive sites or after a membrane-binding step (see appendix). As expected, for the same amount of activated OS, this model demonstrated that the flux for direct recruitment of activated cytosolic proteins, *OS*^*^_(r,t)_, to adsorption sites was higher than that for indirect adsorption through membrane (Fig. 3H).

This model describes that both OS fluxes are characterized by both the rate of transport of activated OS and density of receptors for OS in adsorption sites (Fig. 3) and predict that the probability of phosphorylation for any substrates is dependent of both OS molecular flux and OS residence time. Therefore, controlling the dimensionality of OS oligomers mobility generates different rates of OS flux to adhesive sites and might encode different OS signaling events.

### Different sustained molecular fluxes of OS to adhesive sites encode distinct phenotypic acto-adhesive responses

To test this, we investigated whether the two different fluxes of activated OS could induce different cellular responses. The same frequency of TIRF blue light stimulation was applied on MDCK cells stably co-expressing either OS+GFP-Caax or OS+CIBN-GFP-Caax. Under these conditions, flux of OS oligomers to adhesion sites induced large, curved, auto-assembling centrifugal actin rings after 8 minutes of blue photostimulation (33 mHz, Fig. 4A; Suppl. Video 5). These structures were invadosome rings since characterized by their stereotypical auto-assembling behavior, OS localization and accumulation of LifeAct-iRFP (Fig. 4A) and cortactin (Fig. 4B). Activation of OS presenting a kinase-dead mutation (OS-K295M) or PP2 pretreatment before induction of OS oligomerization abolished invadosomes formation (Fig. S4). By contrast, recruitment of OS oligomers to adhesive sites through membrane diffusion (CIBN-GFP-Caax) did not form invadosomes, but instead induced characteristic lamellipodia (Fig. 4C; Suppl. Video 6). Lamellipodia were highly sensitive to OS activation, as shown by their disappearance only 5 min after stopping photostimulation (kymograph on Fig. 4C). Interestingly, both of these acto-adhesives structures implicated in migration or invasion are poorly present in epithelial MDCK cells. Quantitative analysis confirmed that direct recruitment of OS oligomers to adhesive sites essentially induced invadosome formation but poorly induced lamellipodia, whereas indirect recruitment of OS oligomers through membrane essentially induced lamellipodia and dorsal ruffles (Fig. 4D). Therefore, the two types of OS flux to adhesive sites clearly gave rise to different cellular phenotypes displaying different kinetics. Accordingly, with the same photostimulation frequency, lamellipodia were formed immediately after OS activation (1 min^+/- 0,5 min^), while invadosomes appeared later (15,6 min^+/- 10 min^). To test the relationship between the intensity of each OS flux on the observed cellular responses, we modulated the amplitude of each OS flux by applying light stimulation different frequencies, as previously described^12^. In marked contrast to lamellipodia induction which was apparent for all frequencies tested (from 33 to 2 mHz, Fig. 4F), invadosomes were not able to form if the flux of OS oligomers was not sufficiently intense (by photostimulation greater than a 2 mHz, Fig. 4E).

**Figure 4.**
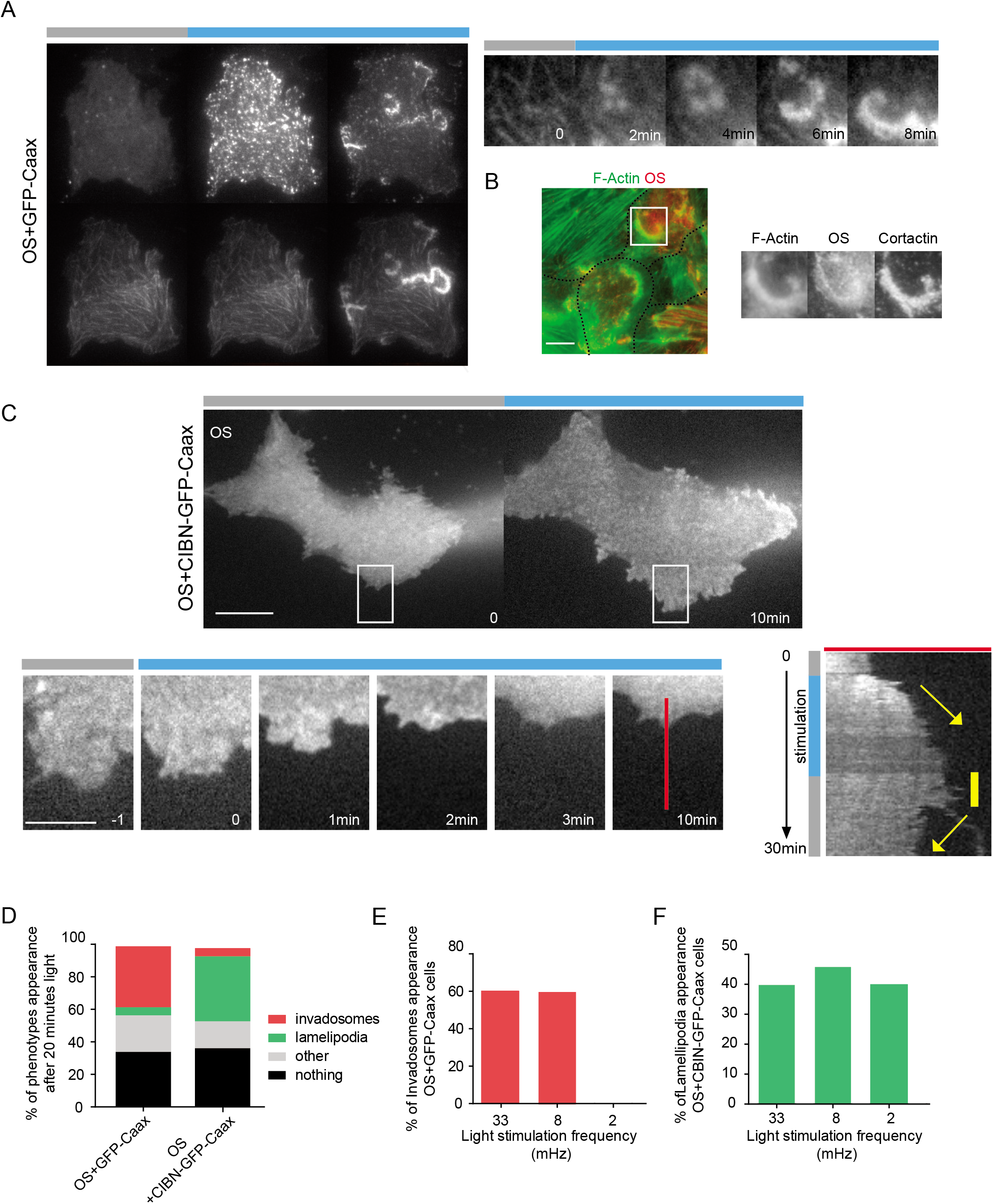
Different rates of OS molecular flux into adhesive sites generate either lamellipodia or invadosomes. **A.** Representative TIRF time serie illustrating how the flux of OS oligomers in adhesive sites induces dynamic invadosome rings (LifeAct-iRFP) in MDCK cells. **B.** These OS-dependent F-actin rings (phalloidin) accumulates the invadosome marker cortactin (confocal imaging). **C.** Representative TIRF time serie illustrating how the flux of membrane-associated OS oligomers in adhesive sites induces lamellipodia in MDCK cells. Kymograph analysis showed high dependency of lamellipodia to blue light (zoom, white square). **D.** Quantification of the different acto-adhesive phenotypes obtained after 20 minutes of blue light stimulation of MDCK cells stably expressing either CIBN-GFP-Caax or GFP-Caax (N<6; > 100 cells per condition). **E-F.** Quantification of the percentage of cells forming invadosomes (E) or lamellipodia (F) depending of blue photostimulation frequency (N=4; > 30 cells per condition). Scale bars: 5 μm.

Our data showed that a sustained flux of OS in adhesive sites promoted invadosome formation, whereas a poorly sustained OS flux induced lamellipodia structures. In conclusion, we could directly manipulate SRC pleiotropy and encode different cellular phenotypes by modulating SRC oligomers flux to adhesive structures, probably through the emergence of different signaling events.

## Discussion

To investigate the SRC coding activity, we engineered a new photosensitive SRC kinase to get spatio-temporal control of SRC conformational intermediates to adhesive sites. We demonstrated that molecular flux modulations of SRC activity were sufficient to generate different downstream signaling pathways and ultimately trigger different cellular responses.

### Engineering synthetic SRC to understand its complex structure-function relationship

Controlling directly SRC signaling in space and time is challenging. Previous protein engineering-based approaches targeted directly the kinase domain of SRC by regulating its selectivity, folding or allostery^14, 15, 23–25^. Chemogenetic approaches are highly specific but are not suitable to achieve min-scale and reversible control of this kinase at the subcellular level. Coupling allosteric regulation of SRC kinase and the light sensitive domain, Lov2, allowed to generate a photo-inhibitable SRC that is reversible and can present subcellular precision^16^. In order to activate SRC in space and time, we engineered a photoactivable SRC based on nontransforming v-Src mutant deleted of its membrane anchoring domain^26^. In addition to improve the spatial resolutionof CRY2-system^12^, engineering OS revealed new functions of SRC conformational intermediates or domains. Despite being mostly described as a monomer, SRC can dimerize and oligomerize after opening of its conformation^8, 10^. In addition to the SH2 domain mediated regulation for SRC focal adhesion targeting, our synthetic OS revealed that SRC oligomerization is a stabilizing process in adhesive sites mediated by SH3 domains. As a result, the apparent contradiction on the essential role of SH2^27^ or SH3^28, 29^ domains in adhesive sites relocalization can now be explained if one assumes that the level of SRC oligomerization is considered since probably controlling specific partnership with a large repertoire of PRR-containing proteins present in adhesive sites. One hypothesis supported by this property is that controlling SH3 avidity or cooperativity by regulating SRC oligomerization is inducing an increase of their apparent affinity leading to new binding as when a minimal level of multivalency of SH3 domains drives phase separation^30^. In addition to modulate c-SRC adaptor functions, formation of oligomers directly controls also kinase activation. CRY2 light-dimerization of cytosolic OS is sufficient to strongly increase Y416 phosphorylation (Fig. 2), and confirmed the existence of a transphosphorylation process^31^.

By integrating intermediates of a signaling node and getting closer to its characteristic spatio-temporal activation patterns, the synthetic approach of optogenetics is complementary to long-term genetic manipulations for exploring new aspects of molecular dynamics *in vivo*.

### Biological relevance of synthetic OS

Our synthetic approach was based on deconstructing some regulatory elements of SRC and this naturally raises the question of its biological relevance. The fact that SRC mutants containing SH2 or SH3 domain deletions can restore invadosome functions in *c-Src -/-* osteoclasts^3^ supported our assumption that other domains of SRC might compensate for the loss of function of point-mutated SH2 domain. Moreover, despite our synthetic biology strategy, we found that OS activation rapidly induced multiple physiological and classical c-

32

SRC-dependent cellular structures such as adhesive sites and actin cytoskeleton structures. Moreover, OS activation mimics activation of thermo-sensitive SRC mutant^18^ and the effects of chemo-controllable synthetic SRC^19, 24^. In MDCK cells, OS activation even induced invadosome rings, which are typical c-SRC-dependent structures in physiological and pathological conditions since highly perturbed in *c-Src* -/- models^2, 33^. The specificity of OS relocalization in adhesive sites is reinforced by our preliminary results on photoactivated optoYes, another member of the SRC family kinases close functionally, showing its inability to be recruited in adhesive sites in response to light (OD personal communication).

In conclusion, even not mimicking physiogical activation of endogenous SRC, OS stimulation is sufficient to activate numerous and characteristic physiological feature of c-SRC signaling pathways. Thus, this synthetic optogenetics supports a new direction to explore the causal link between SRC molecular dynamics *in vivo* and its associated signaling.

### Control of OS molecular flux supports different SRC signaling transfers

By constraining the SRC structure, our optogenetic design decreased the repertoire of potential SRC conformational intermediates and limited the multiple possible molecular mobility behaviors of this signaling network node, which is otherwise highly versatile^34, 35^. This was essential to precisely control the formation of different fluxes of the same OS oligomers in adhesive sites by globally playing on the dimensionality of their mobility (+/- membrane-based recruitment). Thus, our study is in line with previous reports showing that MAPK or integrin signaling tightly rely on the regulation of molecular mobility through cytosol-to-membrane relocalization or receptors clustering^36, 37^. Besides providing a simple qualitative activation approach, we showed that sustaining over 15 min a minimal flux of activated OS in adhesive sites was essential to induce invadosomes (Fig.4E) but not to induce lamelipodia. Our optogenetics approach is an experimental proof-of-concept of a new direction to better understand the molecular basis of pleiotropic molecule in cell signaling. Future studies will determine the molecular determinants of SRC pleiotropy by integrating how the coexistence of both fluxes of SRC and its potential substrates^38^ at adhesive sites can affect the selection process of specific signaling pathways.

In conclusion, exploration of the OS activation regimen revealed new insights into the causal link between molecular signaling and emergence of cellular phenotypes. This approach paves the way to understand the molecular basis of the pleiotropic coding of other signaling nodes and how oncogenic SRC signaling flux leads to physio-pathological transitions without fully compromising the signaling network.

## Methods

Detailed methods are provided in the supplementary data of this paper and include the following:

KEY RESOURCES TABLE (plasmids, reagents, antibodies)

### EXPERIMENTAL CELL MODELS and CULTURE

Experiments were performed on fibroblasts, SYF cells or epithelial MDCK cells transiently transfected and/or stably transduced by viral strategy.

### OPTOSRC PLASMIDS CONSTRUCTION

In this study, c-SRC will be referred to as the wild-type endogenous protein, while SRC represents c-SRC-like activity induced by c-SRC mutants. All the expression plasmids are listed in supplemental STAR Methods table. All the OptoSrc and mutants plasmid construction were cloned in a Nhe1-Not1 digested pSico backbone amplified by PHUSION high fidelity DNA polymerase (NEB) using Gibson assembly (NEB) following the supplier instruction.

### SINGLE CELL OPTOGENETICS EXPERIMENTS BASED ON LIVE IMAGING, FLUORESCENCE RECOVERY AFTER PHOTOBLEACHING (FRAP) AND BRIGHTNESS ANALYSIS

Live imaging, photostimulation and FRAP were performed with an iMIC inverted microscope (FEI) using time lapse transmission, confocal (spinning disk) and TIRF imaging (63x/1.46 oil Korr M27; camera EMCCD, image acquisition with the LA software). Classically, cells were stimulated by 488nm TIRF excitation every 30 seconds (33mHz). OS basal membrane and adhesion sites recruitments were followed using 561nm TIRF imaging, while vinculin-iRFP or LifeAct-iRFP were monitored with 640nm TIRF imaging. Molecular brightness analysis were performed by Fluorescence Correlation Spectroscopy (FCS) acquisition data and performed with the LSM710-Confocor3 confocal microscope (Carl Zeiss), equipped with the C-Apochromat 40x/1.2 water-immersion objective.

### QUANTIFICATION AND STATISTIC ANALYSIS

Data are judged to be significant when p < 0.05 by unpaired, two-tailed Mann-Whitney test. We denote statistical significance as follows: ns, not significant (i.e., p > 0.05); *p < 0.05; **p < 0.01; ***p < 0.001. Graphs and statistical analyses were generated using Prism 6 (Graphpad) and the boxplots indicate the data distribution of the second and third quartile (box), median (line), mean (filled squares), and 1.53 interquartile range (whiskers).

## Supporting information

Ressource tables

## Acknowledgements

We would acknowledge the benevolence and talent Dr S. Wickström for critical reading of this manuscript. We would also thank Dr. C. Gaggioli, Prof. R. Baron, Dr. M. Coppey, Dr E. Miroshnikova, Prof. A. Delon, Dr. I. Wang, prof. M. Digman and prof. E. Graton for experimental help, critical reading and exciting discussions. This work was funded by JCJC ANR « invadocontrol » program, by LLNC as “Equipe labellisée Ligue 2014 and by FRM as Equipe labellisée FRM 2017. A.K. was funded by “Ligue Nationale contre le Cancer” (LLNC) and “la Fondation pour la Recherche Médicale” (FRM). We thank the support of the discovery platform and informatics group at EDyP. Proteomic experiments were partly supported by the Proteomics French Infrastructure (ANR-10-INBS-08-01 grant) and Labex GRAL (ANR-10-LABX-49-01).

## Authors Contributions

A.K., C.O. and O.D. generated constructs and cell lines, performed and analyzed data in cell biology. A.K. and A.G. developed brightness analysis. C.B., A.Kr. and Y.C analyzed mass-spec data and developed pipe-line analysis. B.F. developed the kinetic model and performed computational simulations and analysis.

A.K. and O.D designed the study, and O.D., E.F., C.A-R., A.K., B.F. and C.B. wrote the paper with significant contributions from all authors.

---

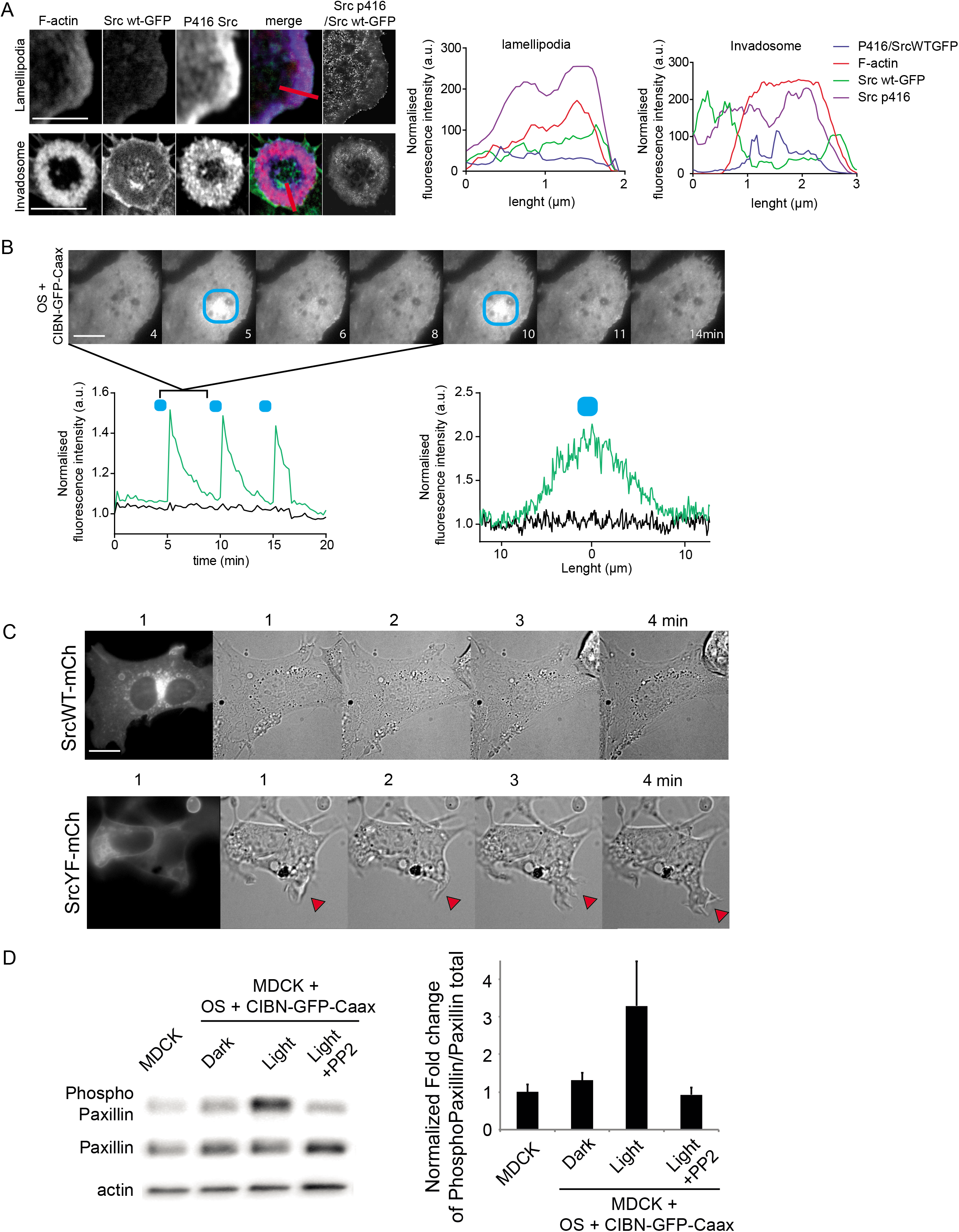

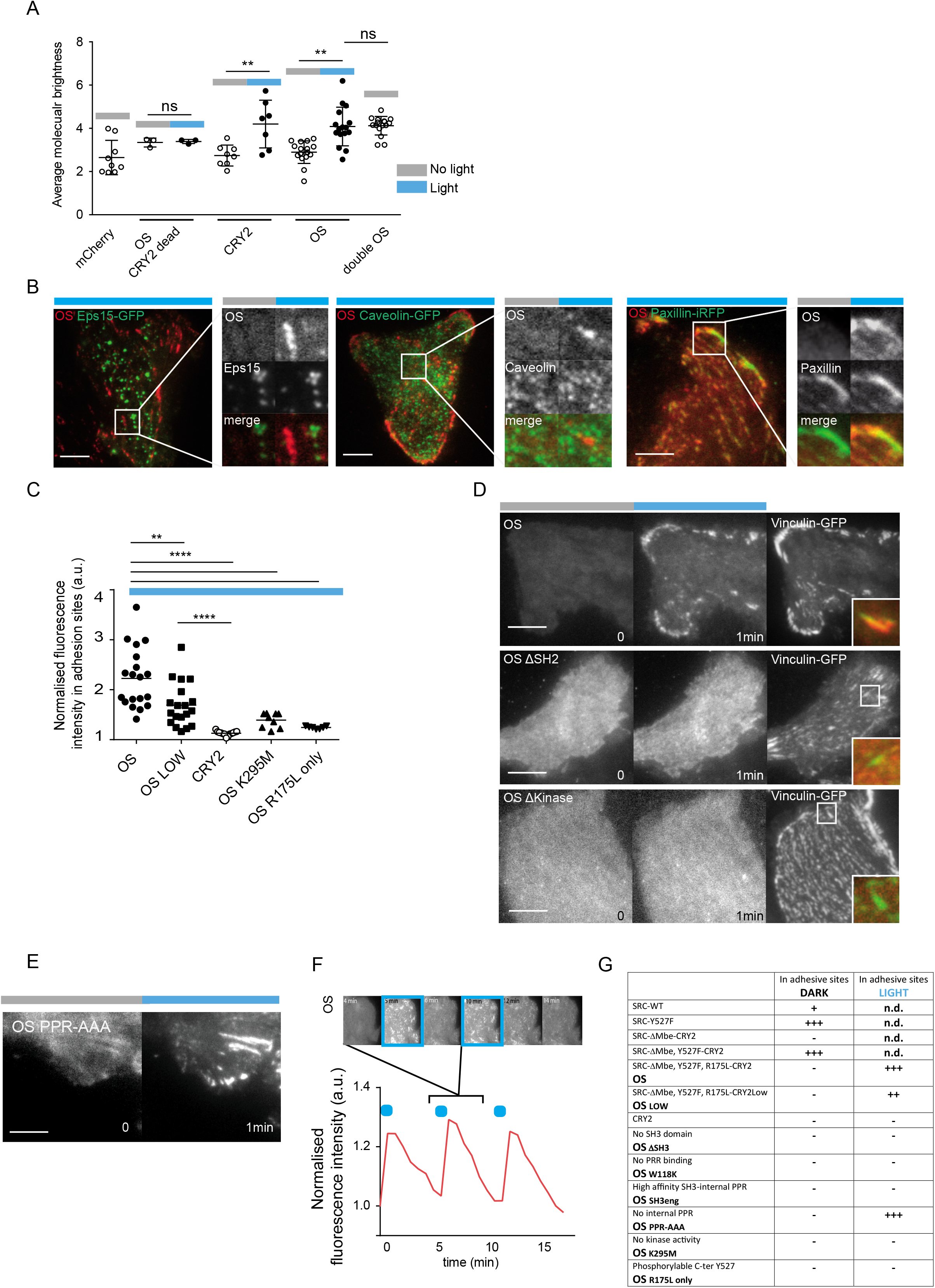

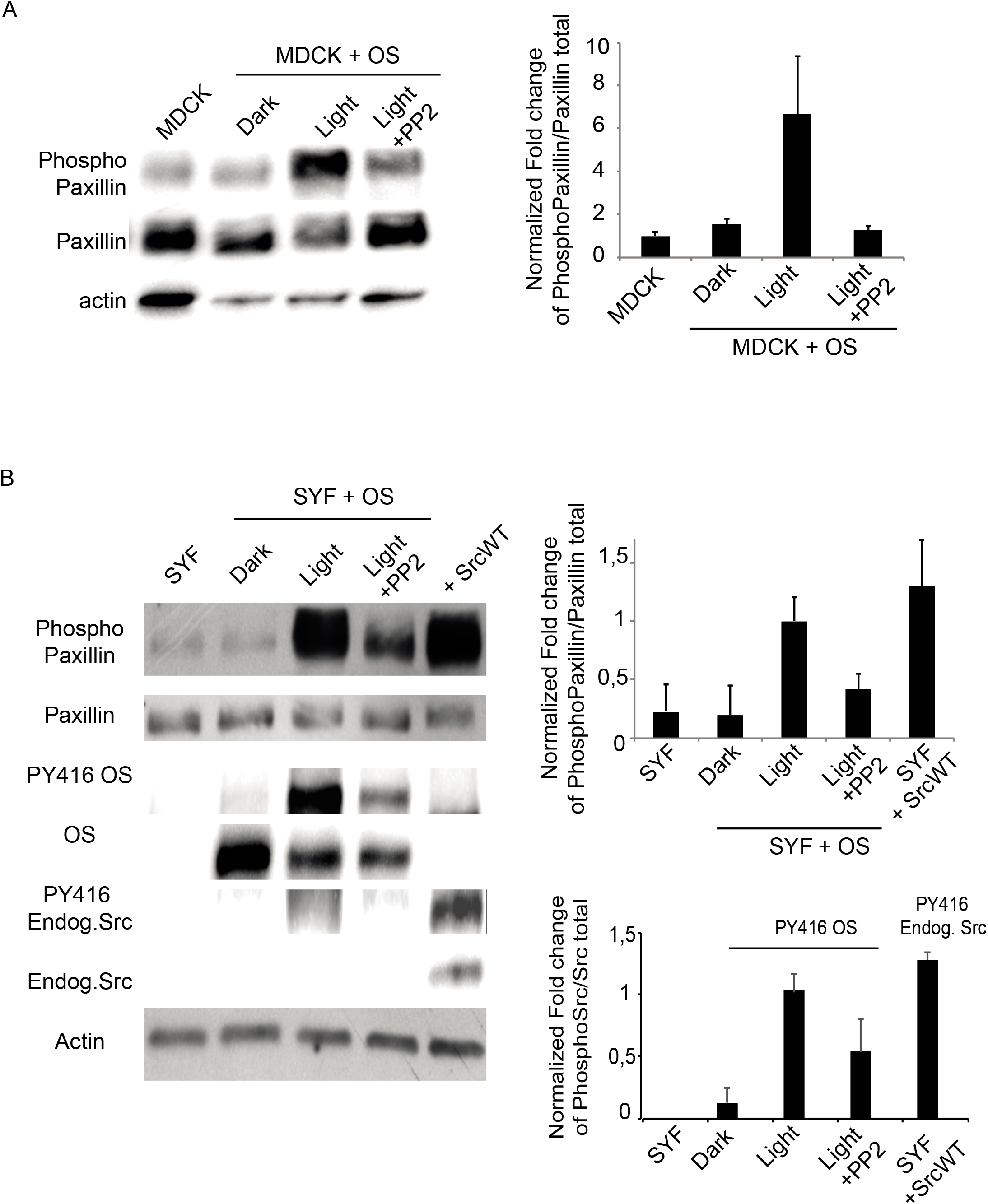

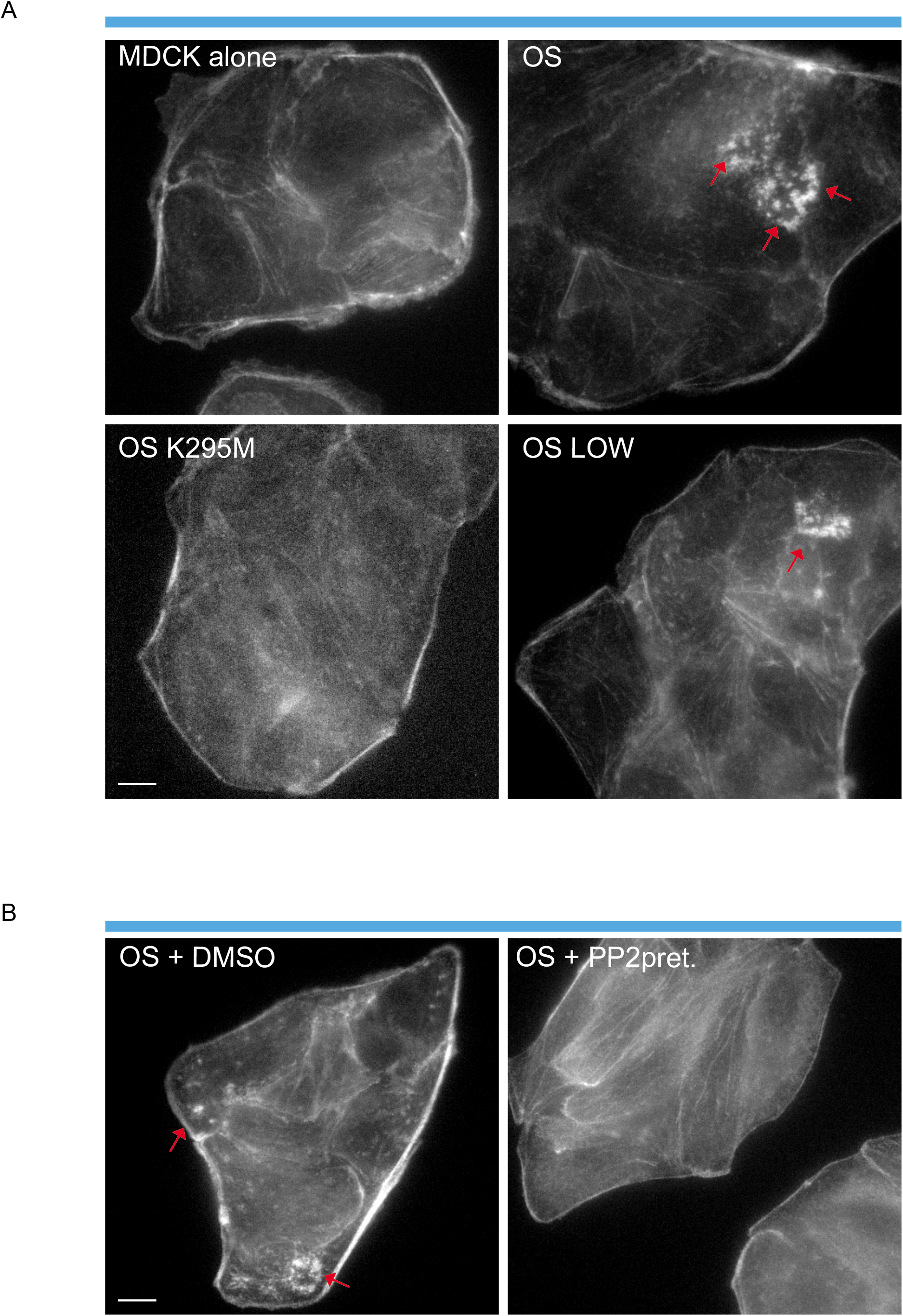

