## Supplementary data and appendix for "Molecular flux control encodes distinct cytoskeletal responses by specifying SRC signaling pathway usage"

***Supplementary information***

*DETAILED STAR * METHODS*

**KEY RESOURCES TABLE** => supplemental Table S3

**EXPERIMENTAL MODEL AND SUBJECTS**

**Cell culture**

All cell lines were cultivated at 37°C and 5% CO2 in DMEM high glucose 4,5g/L, glutamax, PAA) media supplemented with 10% of Fetal bovine serum (GE Healthcare), penicillin and streptomycin 1% (v/v) (PAA).

**METHOD DETAILS**

**Cloning of OptoSrc’s constructs**

In this study, c-SRC will be referred to as the wild-type endogenous protein, while SRC represents c-SRC-like activity induced by c-SRC mutants.

All the expression plasmids are listed in supplemental Table S3. All the OptoSrc and mutants plasmid construction were cloned in a Nhe1-Not1 digested pSico backbone amplified by PHUSION high fidelity DNA polymerase (NEB) using Gibson assembly (NEB) following the supplier instruction.

**Cell Transfection**

All cells were transfected using lipofectamine2000 and according the profocol of the manufacturer (Invitrogen).

**Lentivirus production, cell infection and sorting**

Lentiviruses were produced by co-transfecting pC57GPBEB GagPol MLV, pSUSVSVG and each plasmid of interest using lipofectamine2000 (Invitrogen) in HEK293 FT (precious gift of Dr Nègre from the ANIRA platform) cells plated in 6 well plates at 50% confluency. Media was changed 24hour later. The viral supernatant was collected 72 hours later and filtered with 0.45μm filters. MDCK cells were plated in 6well plates so it achieved 60% confluency the day of infection. The filtered supernatant was directly used to infect cells of interest. The media was changed 24 hours after infection. After 10 days decontamination, cells were FACS sorted (Aria cell sorter 2000, BD) based on the level of expression of mCherry-tagged OS using 561nm LASER.

**Live imaging, OptoSrc photostimulation & recruitment quantification**

Live imaging and photostimulation were performed with an iMIC inverted microscope (FEI) using time lapse transmission, confocal (spinning disk) and TIRF imaging (63x/1.46 oil Korr M27; camera EMCCD, image acquisition with the LA software). Classically, cells were stimulated by 488nm TIRF excitation every 30 seconds (33mHz). OS basal membrane and adhesion sites recruitments were followed using 561nm TIRF imaging, while vinculin-iRFP or LifeAct-iRFP were monitored with 640nm TIRF imaging. OS recruitment were quantified for each TIRF images using time serie analyser plugin on Fiji software and normalized respectively to the first image corresponding to the mCherry cytosolic background (dark), or normalized on vinculin-iRFP/GFP intensity at each time point.

Besides single cell analysis, large cell population photostimulation was perform using a home-made blue LED plate (constructed by C. Tucker, I. Wang and M. Balland, Lihpy, Grenoble-Alpes University). The LED illumination were programmed by Arduino software at 2 minutes frequency at 50% LED power for 5 or 20 minutes.

**Fluorescence recovery after photobleaching (FRAP)**

FRAP analysis was performed on MDCK cell line expressing stably OS +/- CIBN-GFP-caax and OS ΔUD using the iMIC inverted microscope (FEI) and the same 488 nm TIRF protocol (33mHz) to photostimulate OS during 4min before FRAP experiment. Laser photo-bleaching of activated OS in adhesive sites were performed with a 561nm laser (100%) for 150ms (with a bleach area of 10x10 microns). Recovery of activated OS in adhesive site was recorded every 400ms during 1 minute. FRAP analysis were performed on OA offline analysis software using the offline FRAP tool/option (FEI). After normalization on the total cell intensity and on the camera background, a single exponential model was applied A*(1-exp[-t*tau_frap]) in order to measure both characteristic time of recovery and the immobile fraction.

**Brightness analysis of OS oligomeric state**

Molecular brightness can be estimated by analyzing Fluorescence Correlation Spectroscopy (FCS) curves. However, Fluorescence Correlation Spectroscopy (FCS). However, numerous biases and the fit quality of the FCS model can affect this measurement. Thus, we directly estimated the molecular brightness from the raw photon counts over short time intervals in a classical FCS experiment.

Fluorescence Correlation Spectroscopy (FCS) acquisition data was performed with the LSM710-Confocor3 confocal microscope (Carl Zeiss). The inverted AxioObserver stand was equipped with the C-Apochromat 40x/1.2 water-immersion objective, the on-stage cell incubator (PeCon) and the overall environmental chamber stabilized at 37°C at least 1h before acquisitions. Cells were transfected and plated on the 4-well chambered coverslips (LabTek) 24h prior to the experiments at 37° in 5% CO_2_ throughout acquisition. The mCherry fluorescence was excited with the 561 nm DPSS laser, reflected by the 488/561 double primary dichroic filter. The laser light power of 2.6 µW was measured at the objective output at 0.3% AOTF transmission. The fluorescence was selected by the 580 nm long-pass secondary dichroic filter and its fluctuations were sampled by the APD detector at 20 MHz rate. The laser power, the pinhole alignment, and the detected molecular brightness and mobility were carefully controlled on the day-to-day basis using the calibration solution of 15 nM sulforhodamine B (Sigma-Aldrich) in water. The shape parameter (axial-to-lateral size ratio) of the confocal volume of 6.9 was determined by fitting the calibration ACF curve to the single-component free diffusion model, and was fixed to this value in the following analyses. In cases of photostimulation of the optogenetic tag CRY2, the 488 nm line of the Ar laser (1% AOTF transmission) was activated simultaneously with 561 nm FCS laser during acquisition. Due to the high mobility of the cytoplasmic pool of proteins, a stationary regimen of photoactivation was then established and maintained during FCS measurements after a short transition period of ~30 s. Single FCS acquisitions were limited to 10 seconds and repeated 10 times in a cytosolic region of the cell. The presence of the second 488 nm laser increased the apparent brightness of mCherry by *ca.* 8% due to additional excitation in the absorption tail of the fluorophore spectrum. Moreover, the AOTF acousto-optic cross-talk was found to slightly modify the transmitted intensity of 561 nm laser in the presence of 488 wavelengths. The obtained brightness values were accordingly corrected for the proper comparison between no-stimulation and stimulation conditions. The molecular brightness estimated from the FCS curve is also biased by the fluorophore photobleaching and the cytosolic depletion during 488 nm photoactivation. In order to minimize these biases and to be less dependent on the fit quality of the FCS model, we estimated the molecular brightness from the raw photon counts over short time intervals according to the formula:

$$B=\left[ \frac{\sigma^{2}}{<k>-1} \right]\frac{1}{\Delta t}$$

where <k> is the mean number of counts per bin time Δt and σ^2^ is the variance of photon counts. The bin time was chosen to be 150 µs since it is long enough in order to average out the photoblinking effects and to obtain higher value of counts per molecule (CPM), but is still considerably shorter than the average diffusion time of the studied molecules, in order to validate the “pseudo static” approximation. The 1 s integration intervals produce sufficient photon statistics for a good signal-to-noise ratio, and are short enough in order to reduce the effects of photobleaching, cytosolic depletion and cell movements. The binary raw data were saved during the standard FCS acquisitions and were then rebinned and analyzed by a custom written ImageJ macro. The molecular brightness within each cell is averaged from the 60 points (the firsts 40 points are excluded due to the equilibrated state of the beginning of the photostimulation). Each average brightness is then averaged on the +/- 15 cells studied in each condition.

**QUANTIFICATION & STATISTICAL ANALYSIS**

Statistical parameters including the number of n, definition of center and spread (mean ± SD or mean ± SEM), the type of statistical test and correction, and statistical significance are reported in the Figures as well as Legends. Briefly, data are judged to be significant when p < 0.05 by unpaired, two-tailed Mann-Whitney test. We denote statistical significance as follows: ns, not significant (i.e., p > 0.05); *p < 0.05; **p < 0.01; ***p < 0.001. Graphs and statistical analyses were generated using Prism 6 (Graphpad) and the boxplots indicate the data distribution of the second and third quartile (box), median (line), mean (filled squares), and 1.53 interquartile range (whiskers).

*LEGENDS OF SUPPLEMENTARY FIGURES*

**Figure Sup1. Dynamic properties of plasma membrane recruitment of activated OS.**

**A.** Immunofluorescence images of total SRC WT-GFP and its activation marker (PhosphoY416) show the high compartmentalization of activated SRC (PhosphoY416/SRC wt-GFP) in dynamic acto-adhesive structures, such as lamellipodia and the invadosome.

**B.** Representative time series of TIRF images of OS co-expressed with CIBN-GFP-Caax in response to local blue light stimulation. Transient stimulation induces a characteristic rapid (5 sec) and transient (after 300 sec) recruitment of OS to the plasma membrane (blue curve) in comparison with non-stimulated regions (black curve). Local stimulation of OS (co-expressed CIBN-GFP-Caax) presents a minimal spatial resolution of 5 μm around stimulation spot.

**C.** Evaluating the phospho-paxillin/paxillin ratio by western blot and its quantification (SD; N=3) shows the poor leakiness of non-stimulated OS in the dark (stably co-expressed with CIBN-GFP-Caax in MDCK cells) in comparison to MDCK not expressing OS.

**D.** Expression of activated mutant SRCY527F-mCh induces dynamic dorsal ruffles (red arrow) in fibroblasts.

Scale bars: 1 μm (A), 5 μm (C).

**Figure Sup2. Characteristic of OS oligomers relocalization in adhesive sites.**

**A.** Quantification of the average molecular brightness revealed the oligomerization state of OS and OS mutants in response to photoactivation (SD; N=3; >15 cells per condition, unpaired t test).

**B.** Representative TIRF images of activated OS and different markers of subcellular structures, such as clathrin pits (EPS15-GFP), caveolae (caveolin1-GFP) or adhesive sites (paxillin-GFP).

**C.** Quantification of OS, OS LOW (CRY2 mutant presenting decreased oligomerization) and its kinase or C-ter mutants at the plasma membrane after blue TIRF photostimulation (SD; N=3; >30 cells per condition, unpaired t test).

**D.** Representative time series of TIRF images of OS mutants with the SH2 domain or kinase domain deleted and poorly relocalized in adhesive sites (vinculin-iRFP) in response to blue TIRF photostimulation.

**E.** Representative time series of TIRF images of OS PRR-AAA mutants with a non-functionnal internal PRR that is relocalized in characteristic adhesive sites in response to blue TIRF photostimulation.

**F.** Representative time series of TIRF images of OS dimer dynamics in response to blue TIRF photostimulation. Transient stimulation induces rapid and reversible recruitment of OS oligomers to adhesive sites.

**G.** Table summarizing the property of adhesive sites relocalization of the different used mutants.

Scale bars: 2 μm (B, E), 5 μm (D).

**Figure Sup3. Evaluation of OS leakiness and the dynamic range response in different regimens of OS activation and cell types.**

**A.** Evaluating phospho-paxillin/paxillin ratio by western blot and its quantification (SD; N=3) shows the poor leakiness of non-stimulated OS when expressed in MDCK cells and maintained in the dark, in comparison to MDCK not expressing OS.

**B.** Evaluating phospho-paxillin/paxillin and phospho-Y416 (OS or endogenous SRC)/(total OS or endogenous SRC) ratios by western blot and its quantification (SD; N=3) shows the poor leakiness of non-stimulated OS in the dark and expressed in SYF cells (SRC^-/-^ Yes^-/-^Fyn^-/-^ cells). Light-dependent OS dimers induce the same dynamic range of paxillin phosphorylation and kinase activation as did re-expressing­ SRC WT in SYF cells.

**Figure Sup4. Perturbation of the kinase activity of OS affects invadosome formation.**

**A.** Representative confocal images of invadosomes (phalloidin, red arrows) in MDCK stimulated with blue light for 15 minutes and expressing either nothing, OS, OS-K295M (kinase dead) or OS LOW.

**B.** Representative confocal images MDCK expressing OS and stimulated with blue light for 15 minutes. SRC inhibition by PP2 treatment abolished the formation of invadosomes (phalloidin, red arrow) in response to blue light.

Scale bar: 10 μm.

**Supplementary Video 1. Local activation of OS induces a characteristic local SRC-dependent structure, a dorsal ruffle.**

Representative time series (min:sec) of local activation of OS (blue square, 16 mHz) in MEF cells.

**Supplementary Video 2. Light-dependent OS dimerization induces a rapid and specific OS flux targeting adhesive sites.**

Representative time series (min:sec) of blue light stimulation (33 mHz) of MDCK cell expressing OS and vinculin-GFP.

**Supplementary Video 3. Relocalization of OS dimers to adhesive sites is dependent on functional SRC SH3 domains.**

Representative time series (min:sec) of blue light stimulation (33 mHz) of MDCK cell expressing OSΔSH3 and vinculin-GFP.

**Supplementary Video 4. Membrane relocalization of activated OS dimers induces a rapid OS flux targeting adhesive sites.**

Representative time series (min:sec) of blue light stimulation (33 mHz) of MDCK cell expressing OS +CIBN-GFP-Caax and vinculin-iRFP.

**Supplementary Video 5. Induction of OS dimer flux into adhesive sites induces multiple and dynamic invadosome rings.**

Representative time series (min:sec) of blue light stimulation (33 mHz) of MDCK cell expressing OS and LifeAct-iRFP.

**Supplementary Video 6. Membrane relocalization of activated OS dimers induces characteristic large lamellipodia.**

Representative time series (min:sec) of blue light stimulation (33 mHz) of MDCK cell expressing OS+CIBN-GFP-Caax and LifeAct-iRFP.

*Appendix*

In order to specify the flux of OptoSrc (OS) recruitment following light activation, we consider the passive transport mechanisms of activated OS (OS dimers) in the cell volume either with adsorption on the cell membrane or on discrete adhesive sites (Fig.4G). From the mathematical point of view, the model is valid in 2 or 3 spatial dimensions, but it is more tractable in 2D from the numerical point of view. The principal result reported in this Appendix is illustrated in Fig. 4H, where it is demonstrated that the OS flux is sustained at a higher level and on a longer time when activated OS is directly recruited to adhesive sites without intermediate membrane adsorption. Explicit solutions for this problem are given to characterize the different time scales.

Call
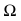
the volume of a spherical cell of radius
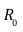
in arbitrary dimensions with boundary
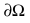
.
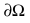
is either a circle in 2D or a sphere in 3D with the same radius. To compute the flux
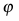
of activated OS per unit length or surface, consider an initial concentration
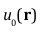
 of light activated OS in the bulk where
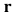
 is the radial vector distance to the origin. The model takes into account the diffusion of OS in the bulk with diffusion coefficient
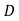
 and introduces an effective adsorption rate
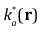
on the boundary
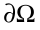
. This rate of adsorption is proportional to the density
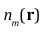
of adsorption sites on
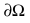
. In presence of CIBN-GFP-Caax,
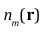
is homogeneously distributed on the membrane and
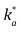
is constant. In contrast,
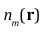
is exclusively concentrated on the adhesive sites in the GFP-Caax case, see Fig. 1 and 4H, and
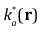
 varies on the boundary
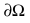

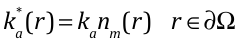

where
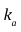
 is the rate per unit surface at full coverage. To compare case 1 where activated OS is recruited to the membrane to case 2, where OS is only recruited to clusters, we assume that $n_{m}(\theta)$ has a square wave distribution, with $n_{m}(\theta)$ alternating between $0$ and $n_{m}$ along the boundary, see App. Fig.1. Following illumination, the two geometries give different flux densities. For the same buffer of cytosolic activated proteins, the flux through adhesive clusters in case $2$ must be larger than in case $1$, since the same amount of material must be adsorbed at infinite time, but on a smaller spatial domain. Using this sink analogy, we also see full adsorption takes a longer time on clusters than on a homogenous boundary.

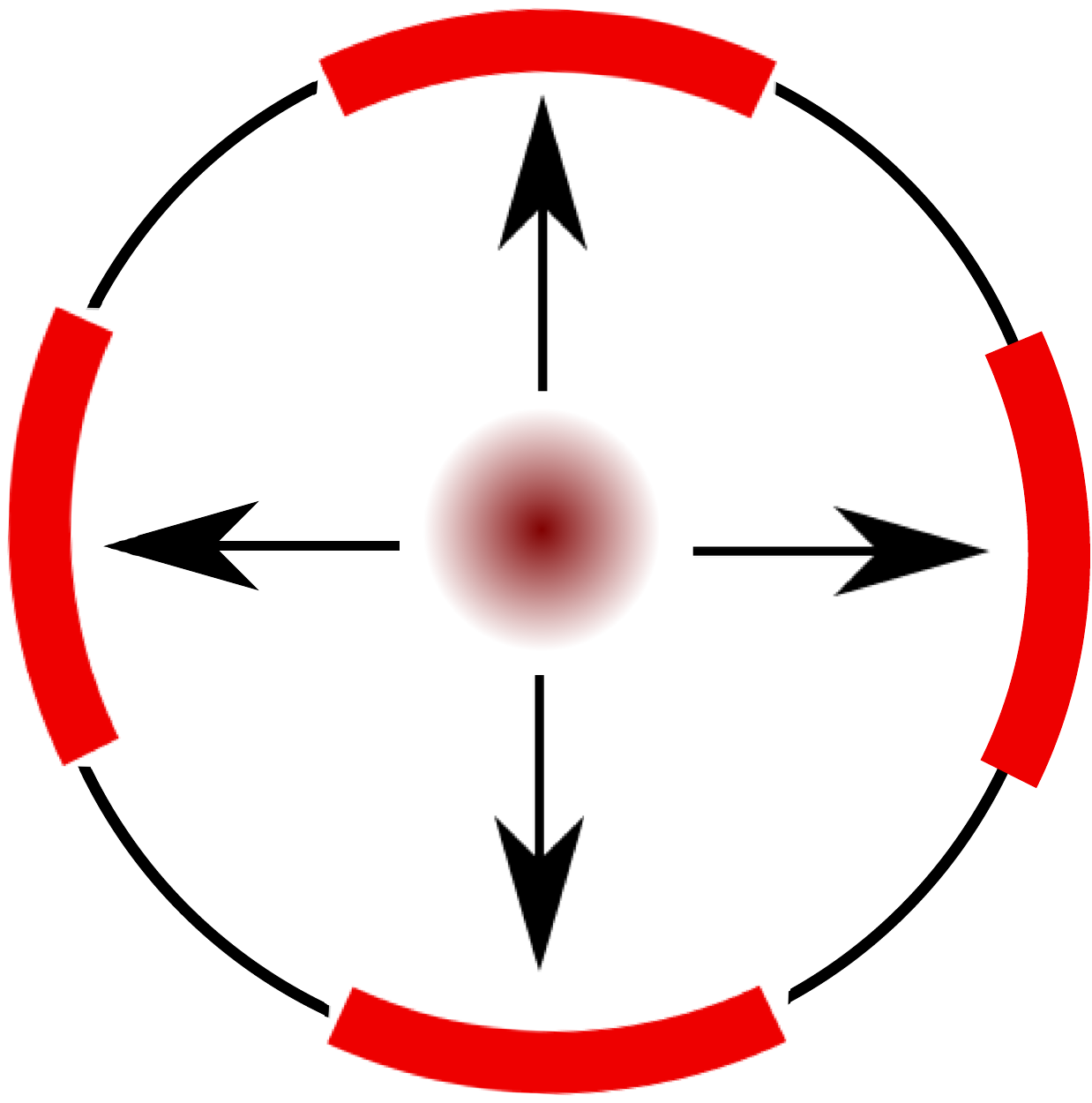

App. Fig.1: Schematic diagram representing a square wave distribution of adsorption sites along the circular domain with $4$ clusters, see case $2$. Clusters of adsorption sites are represented in red. In this case, no adsorption occurs between the clusters. This geometry is in marked contrast with a uniform distribution of adsorption sites on the boundary, see case 1, where proteins can be adsorbed along the perimeter.

To summarize, the concentration $u(r,t)$ solves the following problem:

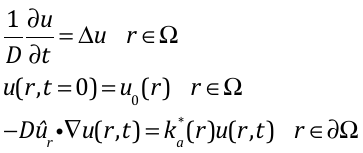

where the first two equations are the diffusion equation together with the initial condition at zero time. The last equation equals the flux φ of adsorption normal to the boundary with the probability that the protein is absorbed on a site assuming that they are in contact (Robin boundary condition). We are interested in computing the time dependent solution of from which the total amount of OS can be calculated.

For what follows, it is useful first to consider the dimensionless ratio to compare the diffusion-limited rate of Schmoluchowski with the rate of OS association with the membrane assuming close contact.

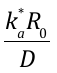

When this ratio is smaller than $1$, we are in the fast diffusion regime where, following illumination, the concentration in the bulk becomes first homogenous and OS proteins subsequently adsorbed. This is the relevant experimental regime(Kaizu et al., 2014),(Northrup and Erickson, 1992) which ensures that our results will not depend on the geometry. Following initial bulk homogenization, the buffer of cytosolic proteins is cleared on a much longer time scale. Calculations below show that this occurs on a characteristic time scale
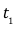
 we compare with the diffusion time scale
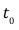

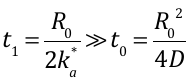

so that the flux
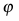
exhibits a sharp increase at short times,
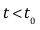
, and decreases more slowly at longer times,
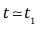
: See Fig. 2. For a typical cytosolic protein with
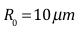
and
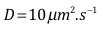
,
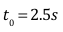
. Since typical association rates of proteins are of the order of a tenth of
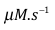
,
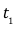
 is of the order of a few ten of seconds. In problem , we have neglected desorption. This approximation is valid if the characteristic time for this process is longer than all other characteristic time and it has the advantage to decouple the adsorption problem from the diffusion problem of adsorbed proteins along the boundary. If diffusion along the boundary occurs, desorption will take place homogeneously along the boundary with small modifications of the bulk diffusion flux contour lines, so that problem is a good approximation at short time.

**In the homogenous case** $\boldsymbol{1}$ (indirect relocalization of activated OS in discrete adhesive sites through a membrane step as in the conditiokn of expression of CIBN-GFP-Caax), our problem is solved in 2D by separation of variables $u(r,t)=\exp[-D\lambda_{m}^{2}t]u(r)$. Sturm-Liouville theory gives the solution as a series

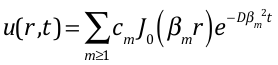

The boundary condition in gives the set
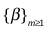
is the ordered set of positive roots of the equation

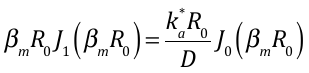

where
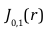
are standard Bessel functions. The coefficients $c_{m}$ are obtained by operating on the initial condition

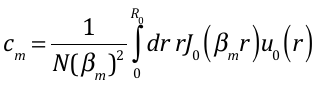

with

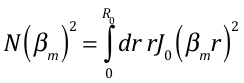

The limit of small $k_{a}^{*}R_{0}/D$ is interesting. In this limit,

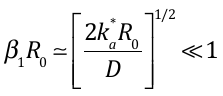

Thus, according to , the characteristic time scale for the exponential decrease in flux density is

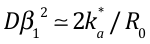

We conclude that the first term in the series dominates and that the flux of proteins adsorbed on the membrane will decrease with a characteristic time $t_{1}=R_{0}/2k_{a}^{\star}$ after a sudden increase on time scale $t_{0}$. The same result holds for the 3D solution with the same symmetry of revolution. Eq. should be adapted using the appropriate basis. Because of dimensional analysis, the same characteristic time scales enters into the problem.

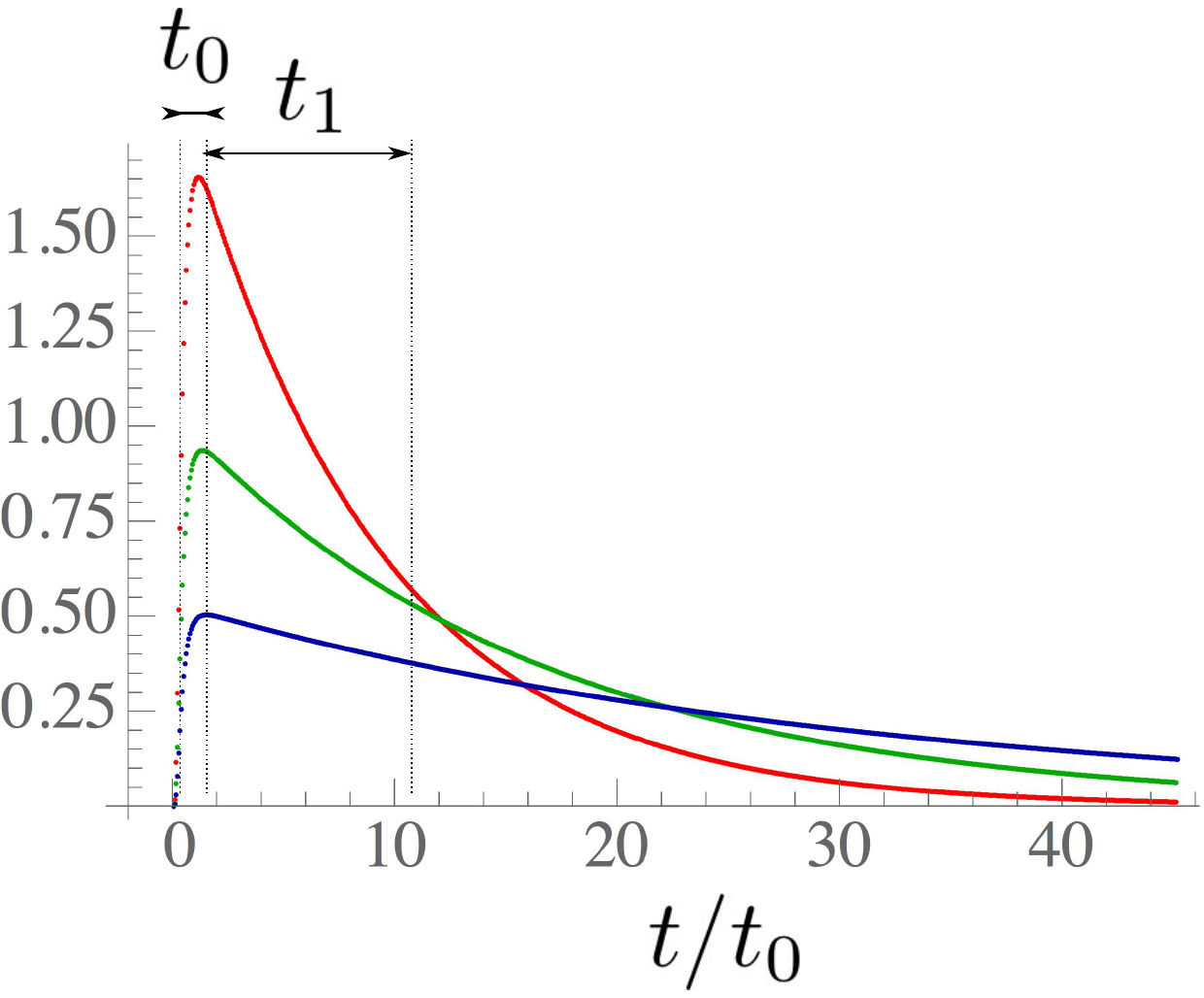

App. Fig.2: Three solutions of the initial problem for the adsorbing mean flux $k_{a}^{\star}u(R_{0},t)$ as a function of $t/t_{0}$ for different values of the adsorption rates $k_{a}^{\star}$ (the bottom curve corresponds to the smallest adsorption rate) All three solutions have the same geometry with $4$ clusters covering $25\%$ of the perimeter. The initial condition is the same in the three cases, so that the area under the curve is the same. The maximum at short time scales with the parameter $k_{a}$ ($k_{a}^{\star}=0.05s^{-1}\mu m^{-1}, u_{0}=0.1\mu m^{-2})$ with an initial distribution taken as a centered gaussian of width $1. For a typical cytosolic protein with R_{0}=10 \mu m, D=1\mu m^{2}/s$, $t_{0}$.= 2.5 s).

App.Fig.2 represents the mean flux per unit of length in a discrete adhesive site for $3$ representative solutions obtained by varying the adsorption rate. Renormalization by $k_{a}^{\star}u_{0}$ is chosen such that the middle curve matches asymptotically $1$ at time $t=0$ when the characteristic time for diffusion $t_{0}$ goes to zero by increasing the diffusion constant. For these representative values of the parameters, the characteristic time $t_{1}$ for clearing the buffer is longer than $t_{0}$. Since the same quantity is cleared at time infinity for all the three cases, the integral under the curve is constant which implies that the maximum at short time increases with $k_{a}^{\star}$.

In the non-homogenous case (direct relocalization of activated OS in discrete adhesive sites as in the condition of expression of GFP-Caax), we solve problem using finite element methods for a square wave distribution of clusters such as in App. Fig. 1. For convenience, we take the density of adsorption sites in the square as equal to the density of adsorption sites for the homogenous membrane.

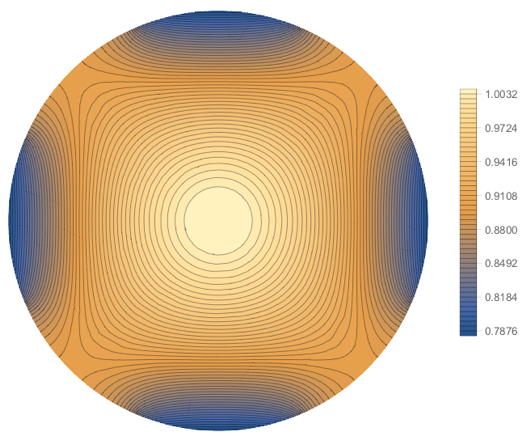

App. Fig. 3. Example of density plot of the OS concentration after illumination for a 2D-solution. The initial condition for the concentration is a Gaussian centered profile normalized to unity. As in Fig.1, the distribution of adsorption sites is a square wave with maximal intensity at the poles and on the equator. The concentration of OS is minimal at these boundaries with a large adsorption flux.

As shown in Fig. 3, the contour lines of the transient solution follow the distribution of adhesive sites with strong gradients near the adsorption boundaries and small gradients between them. App. Fig.4 compares the solution for the homogenous problem with the one for the cluster geometry with averaged quantities per unit length. Since the clusters occupy only $50\%$ of the total perimeter, the area under the curves are the same when conveniently renormalized, so that the total amount of proteins absorbed at infinite time is the same at time infinity. The two curves exhibit the same sharp increase at short time
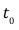
, which only depends on diffusion in the bulk but not on cluster geometry. Again, the mean adsorbed flux is higher in the non-homogenous geometry than in the homogenous case and decreases slowly since all material have to be only adsorbed on the discrete adhesive sites and not between. Depending on the cluster geometry, this happens on different time scales as it can be seen from Eq. by renormalizing the adsorption rate
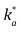
by the fraction of area occupied by the adsorption sites.

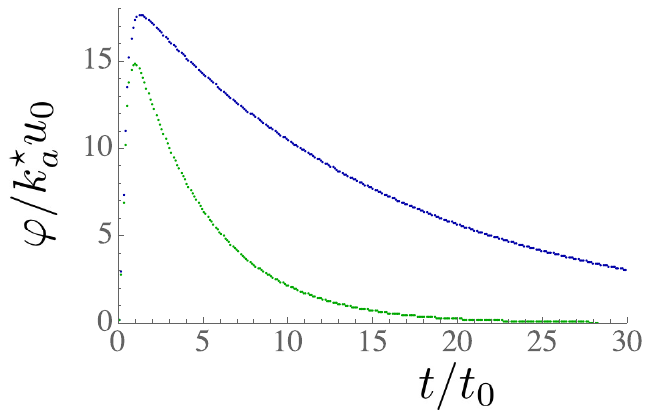

App.Fig.4 Mean flux per unit length in an adsorbing cluster (blue) compared to the solution of the homogenous problems (green). The parameters are the same as in app. Fig. 2.
