## Supplementary material for "Molecular flux control encodes distinct cytoskeletal responses by specifying SRC signaling pathway usage": Ressource tables

| KEY RESOURCES TABLE |  |  |
| --- | --- | --- |
| **REAGENT or RESOURCES** | **SOURCE** | **IDENTIFIER** |
| **Antibodies** |  |  |
| Anti-rabbit Src 32G6 | Cell signaling | #2123 |
| Anti-rabbit phosphoSrc [pY418] | Invitrogen | #44-660G |
| Anti-mouse paxillin 349 | Becton Dickinson | #610052 |
| Anti-rabbit phosphoPaxillin 118 | Invitrogen | #44-722G |
| Anti-mouse p130cas 21/p130 | Becton Dickinson | #610271 |
| Anti-rabbit phospho-p130cas [Y410] | Cell signaling | #4011 |
| Anti-rabbit phospho-p130cas [Y165] | Cell signaling | #4015 |
| Anti-mouse actin AC-40 | Sigma | #5044 |
| Anti-mouse cortactin (p80/85) 4F11 | Upstate | #05-180 |
| IgG2B | Becton Dickinson | #557351 |
| **Chemicals, Peptides, and Recombinant Proteins** |  |  |
| PP2 | Calbiochem | 529576 |
| G sepharose beads fast flow | Becton Dickinson | 55735 |
| NaCl | Euromedex | 1112A |
| NP40 | Fluka | 74385 |
| Sodium desoxycholate | Sigma-Aldrich | D6750 |
| Sodium Fluoride 0.5M | Sigma-Aldrich | 67414 |
| Glycerophasphate | Sigma-Aldrich | G6626 |
| cOmplete EDTA-free protease inhibitor cocktail | Sigma-Aldrich | 11836170001 |
| orthovanadate | Sigma-Aldrich | S6508 |
| TRIS-HCL | Euromedex | 200923-A |
| Glycerol 99.5% | Euromedex | 50405 |
| SDS 20% | Euromedex | EU660 |
| Bromophenol Bleu | Sigma-Aldrich | B0126 |
| OptiMEM | Gibco | 21985-047 |
| Lipofectamine 2000 | Invitrogen | 11668-019 |
| DMEM high glucose | PAA | 31966-21 |
| Fetal bovine serum | Dominique Dutscher | S1810-500 |
| Penicillin streptomycin | PAA | P06-07100 |
| PHUSION | NEB | E0553L |
| Gibson assembly | NEB | E2611L |
| NuPAGE gel | Invitrogen | NP0321BOX |
| **Experimental Models : Cell Lines** |  |  |
| SYF | ATCC | CRL-2459 |
| MDCK | Dr. Isabelle Bonnet (Curie Insitut, Paris France) | N/A |
| MDCK Stable OS+GFP-Caax | This paper | N/A |
| MDCK Stable OS+CIBN-GFP-Caax | This paper | N/A |
| MEF | Home-made | N/A |
| HEK293 FT | Dr Didier Nègre from the ANIRA platform (ENS-Lyon, Lyon France) |  |
| MEF overexpressing SrcY527F | Src transformed MEF Home-made | Petropoulos et. al. 2016 |
| **Recombinant DNA** |  |  |
| cf. recombinant DNA supplementary table |  |  |
| **Software and Algorithms** |  |  |
| FIJI | NIH | <https://fiji.sc/> |
| Prism | Graphpad | <https://www.graphpad.com/scientific-software/prism/> |
| LA | FEI | <https://www.fei.com/> |
| OA | FEI | <https://www.fei.com/> |
| ZEN2010 | ZEISS | <https://www.zeiss.fr/microscopie/telechargements/zen.html> |
| Metamorph | molecular devices | <https://moleculardevices.com/> |
| DIVA (FACS software) | Becton Dickinson | <http://www.bdbiosciences.com/> |
| cytoscape | cytoscape | <http://cytoscape.org/> |
| arduino | arduino | <https://www.arduino.cc/> |
| R | R | https://www.r-project.org/ |
| **Others** |  |  |
| 12 BLUE LED PLATE | Home-made | Irène Wang & Martial Balland (Liphy) |

| **Insert** | **details** | **plamsid** | **References** | **method** | **primers** |
| --- | --- | --- | --- | --- | --- |
| psico | Actin-GFP | pSico | Destaing et. Al. 2010 |  |  |
| GagPol |  | psPAX2 | addgene Plasmid #12260 |  |  |
| VSV-G |  | pCMV | addgene Plasmid #8454 |  |  |
| mCherry | mCherry | pmCherry-N1 | J-L. Coll, Institute for advances biosciences |  |  |
| Cry2 | Cry2PHR - mCherry | pmCherry-N1-cry2 | Addgene Plasmid 26866 |  |  |
| c-Src WT | c-Src(chicken)-mCherry |  | A gift from M. Frame, Beaston Institut for cancer Research, Glasgow |  |  |
| c-Src Y527F | c-SrcY527F-mCherry |  | A gift from M.Frame, Beaston Institut for cancer Research, Glasgow |  |  |
| c-Src Δmyr | c-Src Δmyristoylation(1-13)-Cry2PHR-mCherry | pmCherry-N1 | this paper | Digestion Cry2PHR mCherry par Xho1 | 5'-GCTACCGGACTCAGATgCGGCGCAGCCTGGAGCCACCCGACAGC |
|  |  |  |  |  | 3'-tgtccatcttcatggtggcCGAGCCGGAGCCTAGGTTCTCTCCAGG |
| OS Y527F Δmyr | c-Src Δmyristoylation(1-13) Y527F-Cry2PHR-mCherry | pmCherry-N1 | this paper |  | 5'-GCTACCGGACTCAGATCATGgggagcagcaagagcaagc |
|  |  |  |  |  | 3'-tgtccatcttcatggtggcCGAGCCGGAGCCTAGGTTCTCTCCAGG |
| OS | c-Src Δmyristoylation(1-13) R175L Y527F -Cry2PHR-mCherry | pmCherry-N1 | this paper | by directed mutagenesis with the Quick change II XL site directed mutagenesis kit (Agilent technologies). | 5’ gggaaccgtcagatccgctagccgccaccATGCGGCGCAGCCTGGA |
|  |  |  |  |  | 3’ cgaagttatgcggccgcttacttgtacagctcgtcc |
| OS-LAiR | c-Src Δmyristoylation(1-13) R175L Y527F-Cry2PHR-mCherry-P2A-LifeAct-iRFP | pSico | synthetized cf seq |  |  |
| OS | c-Src Δmyristoylation(1-13) R175L Y527F-Cry2PHR-mCherry | pSico | this paper | 354 amplified pSico diggested by Nhe1/not | 5'-gggaaccgtcagatccgctagccgccaccATGCGGCGCAGCCTGGA |
|  |  |  |  |  | 3'-cgaagttatgcggccgcttacttgtacagctcgtcc |
| OS ΔSH3 | c-Src Δmyristoylation(1-13) R175L ΔSH3(81-142) Y527F-Cry2PHR-mCherry | pSico | this paper | double Gibson | 5'A-gggaaccgtcagatccgctagccgccaccATGCGGCGCAGCCTGGA |
|  |  |  |  |  | 3'A-ccactcttcagcctggatagccagtgccccggcacgctgaggc |
|  |  |  |  |  | 5'B-gggcactggctatccaggctgaagagtggtactttgggaagatc |
|  |  |  |  |  | 3'B-cgaagttatgcggccgcttacttgtacagctcgtcc |
| OS SH3Eng | c-Src Δmyristoylation(1-13) R175L KBRT249-253PPP Y527F-Cry2PHR-mCherry | pSico | this paper | Q5 directed mutagenesis kit | 5'-gggaaccgtcagatccgctagccgccaccATGCGGCGCAGCCTGGA |
|  |  |  |  |  | 3'-cgaagttatgcggccgcttacttgtacagctcgtcc |
| OS ΔSH2 | c-Src Δmyristoylation(1-13) R175L ΔSH2(144-243) Δ(513-532)-Cry2PHR-mCherry | pSico | Construct modified SrcY527F ΔSH2(144-243) construc from Prof. R.-H. Chen … | PCR ampli & gibson (NHE PAC1 in 909) | 5'-gggaaccgtcagatccgctagccgccaccATGCGGCGCAGCCTGGA |
|  |  |  |  |  | 3'-RTTTGTCCATCTTCATGTTAATTAACTGGGGCTCTGTCGAGGTG |
| OS W118K | c-Src Δmyristoylation(1-13) R175L W118K Y527F-Cry2PHR-mCherry | pSico | this paper | by directed mutagenesis with the Quick change II XL site directed mutagenesis kit (Agilent technologies). | 5’-gtcaacaacacggaaggtgacAAGtggctggctcattccctcactacagg; |
|  |  |  |  |  | 3’-cctgtagtgagggaatgagccagccaCTTgtcaccttccgtgttgttgac |
| OS cry2 -dead | c-Src Δmyristoylation(1-13) R175L W118K Y527F-Cry2PHR D387A-mCherry | pSico | this paper | Double gibson | 5'A-gggaaccgtcagatccgctagccgccaccATGCGGCGCAGCCTGGA |
|  |  |  |  |  | 3'A-cacattccaaatcagcaGccaaaagtgtatcccag |
|  |  |  |  |  | 5'B-ctgggatacacttttggCtgctgatttggaatgtg |
|  |  |  |  |  | 3'B-cgaagttatgcggccgcttacttgtacagctcgtcc |
| double OS | c-Src Δmyristoylation(1-13) R175L Y527F-Cry2PHR-mCherry-linker-c-Src Δmyristoylation(1-13) R175L Y527F-Cry2PHR-mCherry | pSico | this paper | 2 étapes 354 ampli : 354+linker in pSico | 5'-gggaaccgtcagatccgctagccgccaccATGCGGCGCAGCCTGGA |
|  |  |  |  |  | 3'-cgaagttatgcggccgcGGTGGCGACCGGTGGATCCCCcttgtacagctcgtccat |
|  |  |  |  | OS-linker + Osrl mCh | 5'-GGGGATCCACCGGTCGCCACCgcggccgcaATGCGGCGCAGCCTGGAG |
|  |  |  |  |  | 3'-tatactatacgaagttatgcggccgcTTACTTGTACAGCTCGTCCATG |
| CIBN-GFP-caax | CIBN(deltaNLS)-GFP-Caax | pSico | This paper (Construct modified from Addgene Plasmid 26867) | ampli of CIBN-GFP-Caax from Plasmid 26867 instered in pSico digested Not1/Nhe1 | 5'-gaaccgtcagatccgctagcATGAATGGAGCTATAGGAG |
|  |  |  |  |  | 3'-tactatacgaagttatgcggccgcCTACATAATTACACACTTTGTCTTTGACTTC |
| GFP-caax |  | pSico | this paper | from pEGFP-C1 |  |
| p130cas-GFP |  |  |  | gift from Maria Fallman, molecular biology UMEA university |  |
| Paxillin-GFP |  | pEF-IRES | Petropoulos et. al. 2016 |  |  |
| Vinculin-GFP |  | pLenti | genebank ID:22330 |  |  |
| caveolin1-GFP |  |  | a gift from J. Lippincott-Shwartz,NIH |  |  |
| eps15-GFP |  |  | a gift from J. Lippincott-Shwartz,NIH |  |  |
| paxillin-iRFP |  | pSico | This paper | Double Gipson amplification | 5'A-ttagtgaaccgtcagatccgATGGACGACCTCGATGCC |
| LifeAct -iRFP |  | pTol2-tetON-IRES | a gift from M.Coppey, Curie Institute |  |  |
| Vinculin -iRFP |  | pTol2-tetON-IRES | a gift from M. Coppey, Curie Institute |  |  |
| LifeAct-GFP |  |  | genebank ID: HQ993061,1 |  | 3'B-tatactatacgaagttatgcTTACTCTTCCATCACGCCG |
